# Comparative analysis of plasma steroid hormone levels reveals lower concentrations in Afrotropical than Palearctic temperate zone passerines

**DOI:** 10.64898/2025.12.13.694095

**Authors:** Ondřej Kauzál, Oldřich Tomášek, Kryštof Horák, Tereza Kauzálová, Judith M. Pouadjeu, Jacques Esembe Chi, Francis Teke Mani, Martin Moos, Zuzana Hochmanová, Zdeněk šimek, Tomáš Albrecht

**Affiliations:** Institute of Vertebrate Biology, Czech Academy of Sciences, Brno, Czech Republic; Department of Ecology, Faculty of Science, Charles University, Prague, Czech Republic; Faculty of Science, Laboratory of Animal Physiology and Phytopharmacology, University of Dschang, Dschang, Cameroon; Bokwaongo Ornithological Club, Buea, Cameroon; Laboratory of Analytical Biochemistry and Metabolomics, Biology Centre of the Czech Academy of Sciences, Branišovská 31, 370 05, České Budějovice, Czech Republic; Department of Applied Chemistry, Faculty of Agriculture and Technology, University of South Bohemia, Studentská 1668, 370 05, České Budějovice, Czech Republic; RECETOX, Faculty of Science, Masaryk University, Brno, Czech Republic; Department of Zoology, Faculty of Science, Charles University, Prague, Czech Republic

**Keywords:** comparative analysis, testosterone, corticosterone, life history, pace of life syndromes, temperate and tropical birds, latitude, stress response

## Abstract

Latitudinal changes in environmental conditions shape the evolution of avian life history traits: tropical bird species typically exhibiting prolonged breeding seasons, a slower pace of life, lower annual investment in reproduction and increased annual survival compared to their temperate counterparts. Major shifts in physiology, reflecting these differences, are expected to be mediated by endocrine regulation, yet comparative data on the sources of inter-specific variation in hormone concentrations remain sparse, especially outside the Nearctic-Neotropic system. We examined variation in two key steroid hormones, corticosterone and testosterone, across 100 passerine species from Europe and the Afrotropics, sampled using standardized field protocols. Baseline levels of both hormones were consistently lower in tropical species, as expected. While stress-induced corticosterone reactivity did not differ between latitudes, it declined with migration distance. Testosterone levels were higher in males than females and declined with migration distance. In tropical species, neither hormone was associated with elevation, another source of environmental variation, but both varied during the breeding cycle, challenging the view of tropical aseasonality. This study broadens the perspective on endocrine correlates of avian life history evolution by testing their generality in an understudied Afrotropical-Palearctic system. The results support the hypothesis that steroid hormone profiles co-evolve with latitude.

## Introduction

Latitudinal gradients offer a valuable framework for studying the evolution of adaptations in related species occupying divergent environments (Schemske *et al*., 2009). Tropical and temperate zones differ markedly in both abiotic conditions and biotic interactions (Schemske *et al*., 2009; Stutchbury & Morton, 2022). Birds have been central to research on latitude-related variation in life history strategies, initially spurred by observations of smaller clutches in tropical birds (Skutch, 1949). Subsequent studies revealed broad-scale differences in life history traits, with tropical species typically exhibiting a slower pace of life, characterised by longer lifespans, higher adult survival, and smaller clutch sizes (Jetz *et al*., 2008; Stevens *et al*., 2013; Valcu *et al*., 2014; Martin *et al*., 2017; Muñoz *et al*., 2018). These life history settings lead to the evolution of a suite of correlated physiological, behavioural and life history adaptations, the pace of life syndromes (Ricklefs & Wikelski 2002). Tropical species typically have lower basal metabolic rates (Wiersma *et al*., 2007a, b; Bushuev *et al*., 2018), reduced baseline blood glucose levels (Tomášek *et al*., 2022), greater resistance to oxidative stress (Jimenez *et al*., 2012), and, in general, greater investment in self-maintenance over reproduction (Ghalambor & Martin, 2000; Ricklefs & Wikelski, 2002) than related temperate species.

Hormones are key regulators of physiological processes in vertebrates, mediating responses to environmental stressors and triggering major life history events such as breeding and migration (Sapolsky *et al*., 2000; Dawson, 2008). Among the most studied avian hormones are corticosterone and testosterone. While both have broadly conserved functions across vertebrates, their optimal circulating levels can vary substantially among and within species (Romero, 2002; Garamszegi *et al*., 2005, 2008; Bókony *et al*., 2009).

Corticosterone (CORT), the primary avian glucocorticoid, governs energy balance, gluconeogenesis, and acute stress responses (Wingfield *et al*., 1998; Sapolsky *et al*., 2000; Millanes *et al*., 2024). Baseline CORT, which supports routine metabolic functions (Sapolsky *et al*., 2000; Jimeno & Verhulst, 2023), is generally lower in Neotropical than in North American temperate species (Hau *et al*., 2010), possibly reflecting their lower basal metabolic rates (Wiersma *et al*., 2007a). Stress-induced CORT increases rapidly in response to acute challenges, redirecting energy allocation toward immediate survival (Sapolsky *et al*., 2000) and has been positively linked to lifespan and survival across species (Hau *et al*., 2010; Taff *et al*., 2022). It is thus hypothesised to be higher in tropical species, where selection may favour enhanced self-maintenance and predator avoidance (Stutchbury & Morton, 2022). However, comparative analyses yielded mixed results: some found no latitudinal trend (Hau *et al*., 2010), others a positive correlation with latitude (Bókony *et al*., 2009; Jessop *et al*., 2013).

Testosterone (TEST) mediates male reproductive behaviours such as territorial defence, song, parental care or gonadal development and spermatogenesis (Wingfield *et al*., 2001). Across species, TEST shows substantial variation. Tropical birds typically have lower overall and peak breeding-season levels compared to temperate species (Goymann *et al*., 2004; Hau *et al*., 2010; but see Goymann *et al*., 2006). These patterns may underpin broader differences in sexual selection intensity between tropical and temperate birds (Albrecht *et al*., 2013; Stutchbury & Morton, 2022; Cooney *et al*., 2022). On top of that, the prevalence of female song and duetting (Tobias *et al*., 2016; Mikula *et al*., 2019) and the greater involvement of females in territory defence in tropical species (Barber *et al*., 2024) may lead to more similar TEST profiles between sexes. Both CORT and TEST, when chronically elevated, can supress immune functions (Foo *et al*., 2017; Sur *et al*., 2025), potentially reducing survival (Bicudo *et al*., 2010; Maness *et al*., 2023). Therefore, low concentrations of both hormones or greater sensitivity to their changes (Sapolsky *et al*., 2000; Canoine *et al*., 2007), may represent adaptations to longer breeding seasons or life histories prioritising survival over reproductive investment in the tropics (Stutchbury & Morton, 2022). On the other hand, even species living in relatively benign tropical environment show seasonal variation in hormone levels (Chiver *et al*., 2014; Lopes *et al*., 2021; Gonzalez-Gomez *et al*., 2023), further highlighting the need to account for such variability in comparative analyses.

Hormone profiles are also shaped by migratory behaviour (Krause *et al*., 2021; Bauer & Watts, 2021; Gonzalez-Gomez *et al*., 2023). Importantly, many of long-distance migratory species breed predominantly in northern temperate regions, rather than in the tropics. Energy mobilisation facilitated by CORT plays a key role during the preparation for migration and during the migration itself (Gonzalez-Gomez *et al*., 2023), though breeding season CORT appears unrelated to migratory status across species (Uehling *et al*., 2024). Long-distance migrants generally have higher survival and smaller clutches compared to sedentary species (Bruderer & Salewski, 2009; Winger & Pegan, 2021), suggesting a slower pace of life and therefore lower TEST (Hau *et al*., 2010). Therefore, it is noteworthy that the only study investigating TEST and migration reported higher TEST in migrants than sedentary species (Garamszegi *et al*., 2008), possibly reflecting their shorter, more synchronous breeding seasons (Spottiswoode & Møller, 2004).

Another ecological factor, that may confound latitudinal comparisons between tropical and temperate species, is diet. Species with sugars rich diets, such as nectarivores and frugivores, are more common in the tropics (Stutchbury & Morton, 2022), and may exhibit higher blood glucose levels than other birds (Tomášek *et al*., 2022; but see Szarka & Lendvai, 2024). Whether this is reflected in levels of CORT, the primary avian glucocorticoid, however, remains to be tested. Elevation gradients represent another major axis of environmental variation shaping avian life histories. Montane species often exhibit distinct life history traits, including higher adult survival and smaller clutch sizes, despite higher metabolic rates (Hille & Cooper, 2015; Boyce *et al*., 2015). In an Afrotropical-European comparison, tropical montane birds showed elevated baseline and stress-induced glucose levels, a key circulating energy substrate, relative to lowland species (Tomášek *et al*., 2022). Whether this is mirrored by variation in CORT remains unknown. Similarly, elevated TEST levels have been reported in high-elevation Neotropical species, possibly reflecting selective pressures associated with shorter breeding seasons and stronger sexual selection, paralleling patterns observed between temperate and tropical zones (Goymann *et al*., 2004).

Most comparative studies of avian hormones, particularly CORT and TEST, have relied on data extracted from literature or global databases (such as the HormoneBase: Vitousek *et al*., 2018) which are often based on uneven or unstandardized sampling (e.g. Goymann *et al*., 2004; Garamszegi *et al*., 2008; Casagrande *et al*., 2018; Husak *et al*., 2021). A notable exception is Hau *et al*. (2010), who directly measured CORT and TEST in wild birds across latitudes. However, their study was limited to a relatively small number of New World taxa, included only males, and did not account for seasonal variation or other potentially confounding factors, neither for within species variability in steroid hormone concentrations. Although the focus on hormones is central to avian comparative ecophysiology, much of our current understanding of the physiological adaptations in tropical birds is still based on Neotropical species (see Goymann *et al*., 2004; Hau *et al*., 2010; Moore *et al*., 2019; but see Goymann *et al*., 2006; Apfelbeck *et al*., 2017). The Afrotropical realm is particularly underrepresented (but see Albrecht *et al*., 2013; Horák *et al*., 2022, in press; Tomášek *et al*., 2022), which severely limits the generalisability of findings across continents. However, global analyses demonstrate consistent life history differences between tropical and temperate species worldwide (Jetz *et al*., 2008; Stutchbury & Morton, 2022), and several studies provide indirect evidence that Afrotropical species also have a slow pace of life, inferred from smaller clutches (Jetz *et al*., 2008), longer lifespans (Stevens *et al*., 2013; Valcu *et al*., 2014), or lower baseline blood glucose levels (Tomášek *et al*., 2022).

Here, we present the first comprehensive field study examining the parallel evolution of circulating steroid hormone levels in phylogenetically related temperate and tropical passerine species from the Palearctic and Afrotropical realms. The study is based on a unique dataset comprising 301 individuals of 100 species from 31 families, sampled following a standardized field protocol. Crucially, our study accounts for multiple ecological and physiological covariates often overlooked in previous research. Building on earlier findings from Nearctic-Neotropical comparisons, as well as glucose concentration data from the Afrotropics (Tomášek *et al*., 2022), we tested the following predictions: (1) Because of their faster pace of life, baseline CORT is higher in temperate zone than tropical species (Hau *et al*., 2010; Apfelbeck *et al*., 2017), and positively associated with clutch size and negatively with body mass (both between and within species; Casagrande *et al*., 2018; Vitousek *et al*., 2019); (2) Stress-induced CORT reactivity is higher in Afrotropical species, where it may support enhanced adults survival (Hau *et al*., 2010); (3) TEST levels are lower in tropical species, which generally experience less intensive competition for mates and have lower levels of sexual promiscuity (Garamszegi *et al*., 2005; Stutchbury & Morton, 2022). TEST is higher in males but the difference between sexes is lower in tropics, where females regularly contribute to territory defence and singing (Tobias *et al*., 2016; Mikula *et al*., 2019). Finally, (4) migratory behaviour may be associated with either higher TEST, reflecting the short and intensive breeding seasons of migrants (Garamszegi *et al*., 2008), or lower TEST, consistent with the slower pace of life of long-distance migrants (Wiersma *et al*., 2007b), whereas no association is predicted for CORT (see Bauer & Watts, 2021). (5) We also predict higher baseline CORT and TEST in montane versus lowland tropical species due to their higher energy demands, shorter breeding season and stronger sexual selection (Goymann *et al*., 2004), and (6) evaluate the effects of dietary contents on CORT. This approach enables a more nuanced understanding of hormone variation across broad environmental gradients and biogeographic realms.

## Materials & methods

### a) Study species and field sites

We trapped small passerines using mist nets during regular fieldworks in Czechia (temperate zone site, 48.69-49.74 °N, 13.94-16.91 °E, 180-730 m a.s.l.) and along a well-established elevation gradient on Mt. Cameroon (4.1 °N, 9.1 °E, 10-2280 m a.s.l.) in 2017-2019 (Supporting Information, Fig. S1). To obtain breeding season values for most species, including the few tropical species breeding in the rainy season, fieldwork was conducted in Cameroon both during the dry season (in November and December) and the rainy season (in August and September), and in Czechia between April and July (Supporting Information, Table S1). Mist netting was conducted in various habitats (from streams, reedbeds, open shrubland to secondary or primary forest and urban areas) in order to cover as ecologically and taxonomically diverse spectrum of passerine species as possible.

### b) Field data & molecular sexing

To obtain the baseline level of both CORT and TEST, we collected small blood sample (maximum of 0.5 % body mass) from 301 individuals of 100 species (44 temperate, 56 tropical) using jugular vein venipuncture within 4 minutes of capturing the individual (range 49-256 s, average 116 s), which is generally considered as a baseline level in literature (Romero & Reed, 2005; Bókony *et al*., 2009; Deviche *et al*., 2016). We then collected another blood sample after 30 min (range 27-37 min, average 30 min) of standardized stressor (keeping the bird in a bird bag) to determine the stress level of CORT. For the analysis of stress-induced CORT, we carefully selected a subset of 75 individuals of 24 species in order to get a balanced dataset across passerine phylogeny with the aim to select closely related species pairs where possible. Blood samples were stored on ice or in fridge upon their centrifugation to separate plasma and red blood cells, both were subsequently transferred to different tubes and deep frozen in liquid nitrogen in the field (range 15-220 min, average 85 min, from the point of collecting the sample to deep freezing it) and then stored in – 80 °C until further analysis.

All birds were ringed with unique metal ring (issued either by Czech Ringing Centre or SAFRING), standard morphological measurements were taken and the breeding status of each bird was assessed (based on brood patch and cloacal protuberance development), followed by the release of the bird. Because brood patch and cloacal protuberance development can be unreliable for sexing in often monomorphic tropical species, all individuals of tropical species were molecularly sexed. DNA was extracted from a small blood sample preserved in 96 % ethanol using DNeasy Blood & Tissue Kit (QIAGEN). Then either a part of the CHD1 gene (Griffiths *et al*., 1998) or AT5A1 gene (Bantock *et al*., 2008) were amplified and the PCR products were examined using gel electrophoresis. In unclear cases, capillary electrophoresis was used to separate the PCR products (Synek *et al*., 2016).

### c) Life history traits

We compiled life history data for each species from literature and other sources, namely clutch size (Myhrvold *et al*., 2015) and diet (Wilman *et al*., 2014). Data on species-specific breeding status, individual baseline blood glucose, and, for tropical species, information on elevational distribution were obtained from our previous work on an overlapping dataset (Tomášek *et al*., 2022). Average body mass for each species was calculated from our own data. We were unable to obtain clutch size information from literature for 5 tropical species, therefore, we used clutch size from their closest relatives (Supporting Information, Table S2).

For each migratory temperate zone species, we calculated the centroid of their non-breeding distribution using QGIS 3.28 (QGIS.org, 2025). The non-breeding distribution was created by uniting “year-round” and “winter” distribution polygons from source data (BirdLife International and Handbook of the Birds of the World, 2018) which were then restricted to the geographical limits of Europe and Africa where all analysed species overwinter (Cepák *et al*., 2008), with the exception of the Common Rosefinch (*Carpodacus erythrinus*) which overwinters in Indian subcontinent (Lisovski *et al*., 2021) and its non-breeding centroid was calculated from non-restricted data. The migration distance was calculated as a circular arch with radius of 6378 km (radius of Earth) and with the angle as the difference between latitude of sampling and latitude of the wintering grounds centroid. For the resident temperate species and for all tropical species the migration distance was set to 0.

To test for possible confounding effect of common ancestry, a phylogenetic tree for the studied species was constructed using publicly available DNA sequences (for details see: Horák *et al*., 2022).

### d) LC-MS/MS analysis

All samples were randomized and then analysed in an analytical chemistry laboratory in RECETOX (Brno, Czechia) using liquid chromatography–tandem mass spectrometry (LC-MS/MS) and a newly developed method utilizing hormone derivatization to increase reaction specificity (Bílková *et al*., 2019) adapted for plasma samples (for detailed description see Supporting Information: Materials and methods).

Out of the 301 samples of circulating plasma TEST measured, 130 were under quantification limit (60 pg/ml). Their distribution was non-random in the dataset but rather followed expected biological distribution (Supporting Information, Table S3).

### e) Statistical analysis

For each individual, the difference of its body mass measured in the field from average species-specific body mass (further referred in text as *individual body mass deviation; IBMD*) was calculated as well as the difference of the altitude of capture and species-specific elevational midpoint. Average body mass and all hormone values were ln transformed. The sex was coded as dummy variable (0: male, 1: female) as well as breeding latitude (0: temperate, 1: tropical zone), breeding season (0: breeding, 1: nonbreeding) and dry/rainy season (0: dry season, 1: rainy season). The information about proportions of various types of diet for each species from source data was merged into three categories: carnivory, herbivory and nectarivory; and then logit transformed (for details see Tomášek *et al*., 2022). Carnivorous and herbivorous diets were strongly negatively correlated (Pearson’s r = – 0.83, p < 0.001), therefore we used only carnivory and nectarivory in the further analyses, such that carnivory can be interpreted as a position of species on the herbivory-carnivory continuum.

Data were analysed using the Bayesian phylogenetic mixed models based on the Hamiltonian Monte Carlo algorithm using the brms package v. 2.22.0 (Bürkner, 2017). Default priors defined in the brms package were used and the models were run in 8 chains, each with 8,000 iterations, warm-up of 4,000 and thinning of 1. All results are presented as posterior means with quantile-based 95% credible intervals (CrI95). The support for an effect is considered significant when CrI95 does not contain zero. Bayesian R^2^ were calculated using the performance package v 0.13.0 (Lüdecke *et al*., 2021) and brms models were compared using the expected log predictive density from leave-one-out cross-validation (“elpd loo”) using the package loo v 2.8.0 (Vehtari et al., 2017). All analyses were performed in R v. 4.4.2 (R Core Team, 2023). Convergence and sampling efficiency of all brms models were assessed by ensuring there were no divergent transitions after warmup, Gelman-Rubin statistic values were ≤ 1.01 and all chains exhibited E-BFMI values greater than 0.7 (Bürkner, 2017; Vehtari *et al*., 2021).

We treated the values for TEST below the quantification limit (“left-censored data”) in four different ways and compared the results which were consistent across all methods (for details of these methods see Supporting Information, Material and methods, for results of the main models see Supporting Information, Table S4). We selected the method using the native *cens* function of the brms package for treating left-censored data for all further models reported in the main text as, unlike the rest of the methods used, it also accounts for the uncertainty of the imputed values (Bürkner, 2017).

Apart from the potentially confounding effect of phylogeny, all analyses were controlled for the species average body mass and the IBMD. We have found significant positive effect of sampling time for both hormones and negative effect of time since sunrise for CORT (Supporting Information: Materials and Methods and Table S5). Therefore, all further analyses were controlled for these two variables. There was no effect of playback in the data (supplementary Materials and methods, table S6. Apart from being included in the main models, all predictors were also tested individually in models hereafter referred to as “adjusted single effect” (ASE) models to test their overall effects. All the ASE models were controlled for the species average body mass, IBMD, sampling time, time since sunrise and sex (supplementary Materials and methods, Table S7).

## Results

Values of baseline CORT were available for 301 individuals representing 100 passerine species – 130 samples of 44 temperate species and 171 samples of 56 tropical species – average 3.0 samples/species (s.d. = 1.55, Fig. 1). Same individuals were analysed also for TEST. Stress-induced levels of CORT were measured in a subset of 45 ind. of 13 tropical species and 30 ind. of 11 temperate species. Repeatabilities of the within species hormone measurements ranged between 0.15 and 0.29 and were all statistically significant (Table 1).

**Figure 1:**
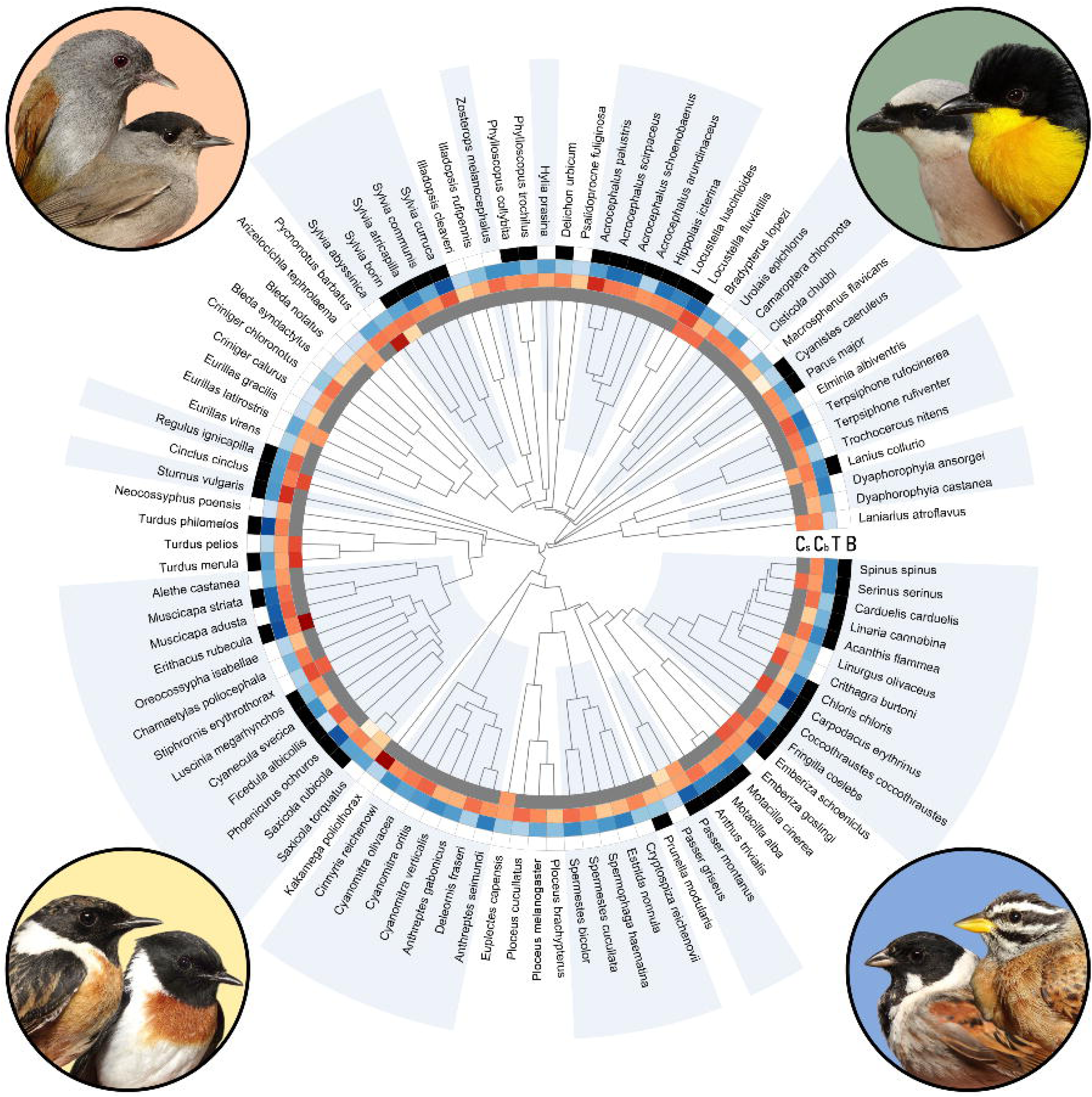
Distribution of mean values of stress corticosterone (C_s_), baseline corticosterone (C_b_), testosterone (T) and breeding latitude (B) across study species (*n_spec_* = 100). Darker shades mean higher values of hormones, grey colour means no data, black are temperate zone species and white tropical ones. Taxonomic families are separated by alternate background shading.

**Table 1:**
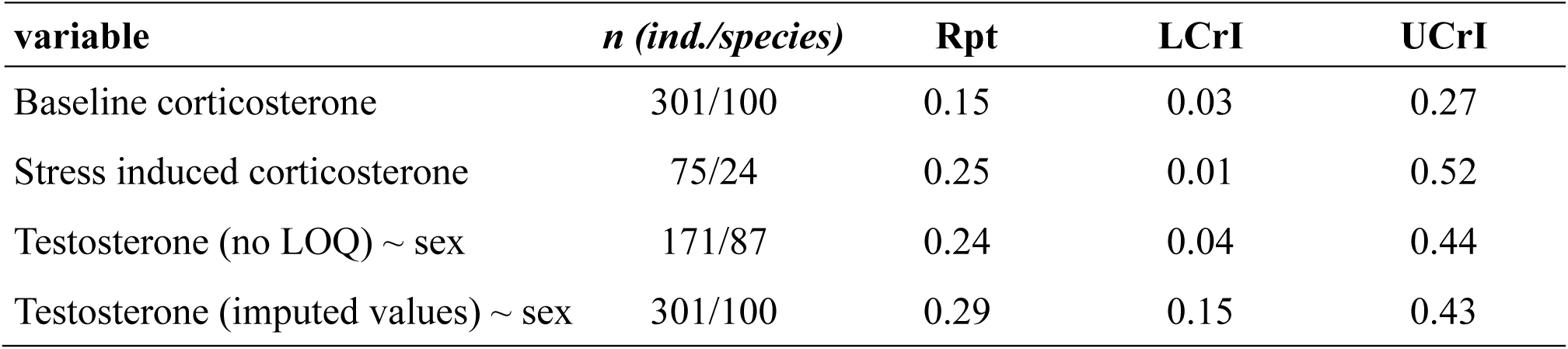
Sample sizes and within species repeatabilities and their respective 95% credible intervals. Because of great differences of TEST between males and females, repeatabilities of TEST were also controlled for sex.

### a) Corticosterone

We first tested whether baseline CORT levels vary with breeding latitude and whether this variation differ between sexes using a model without and with an interaction between sex and breeding latitude. In both models, tropical species showed lower baseline CORT in general. The interaction term in the more complex model indicated that the difference in CORT between tropical and temperate species was primarily driven by males (Fig. 2a). There was some support, albeit not decisive, for the interaction term (95% CrI: –0.03, 0.83; *Δelpd loo* = 0.6, s.d. = 2.1; Table 2), therefore we decided to present the model with the interaction as the main model to retain this potentially interesting source of variation (Fig. 2a). Full results of the selected model are shown in Table 2, the alternative model is presented in Supporting Information, Table S8 and results of ASE models in Supporting Information, Table S7.

**Figure 2:**
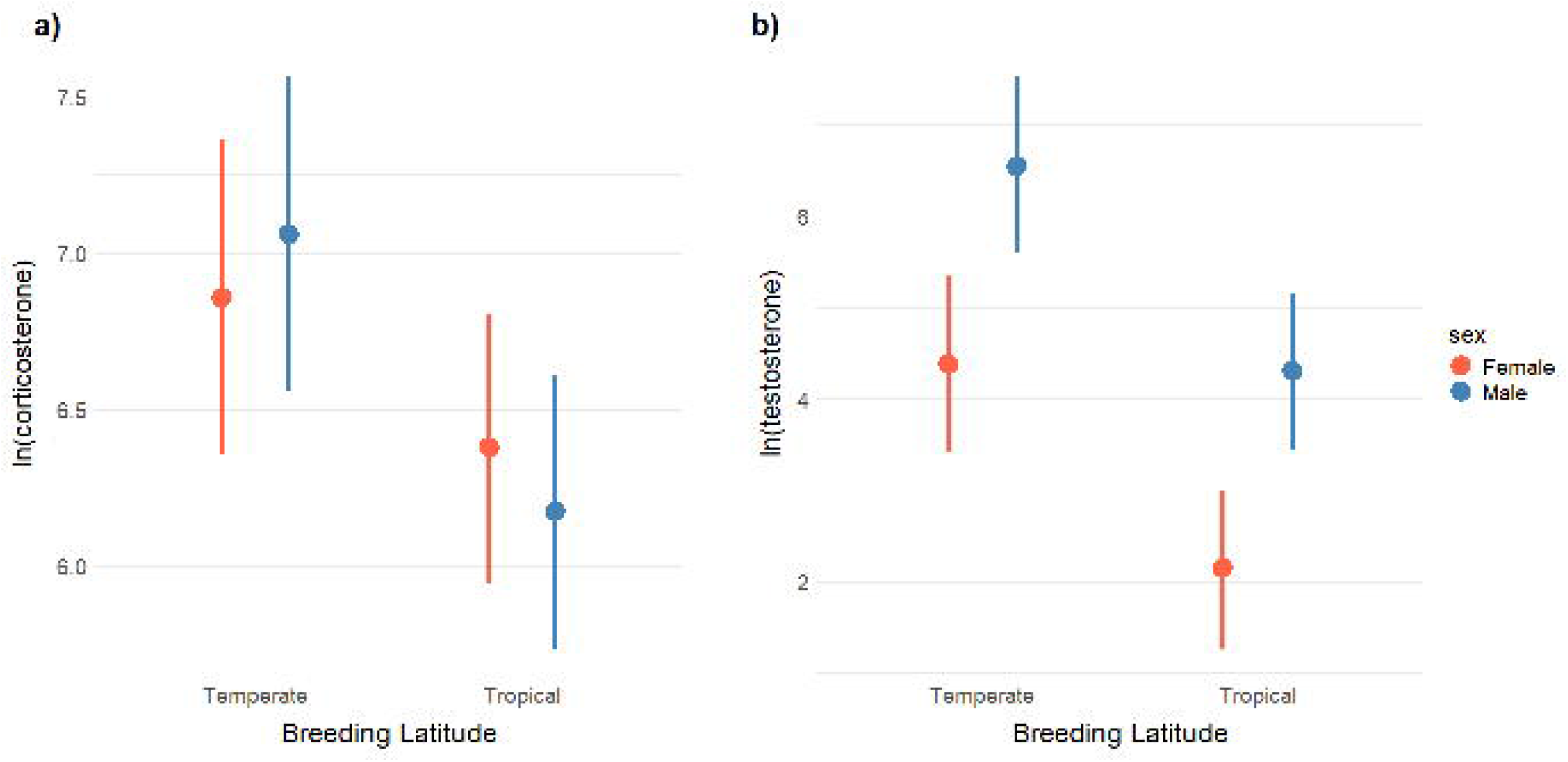
Posterior predictions of a) ln transformed baseline CORT and b) ln transformed TEST for both sexes and breeding latitudes based on the respective Bayesian main models (Tables 2 and 3). Points represent posterior median predictions and vertical bars show 95% credible intervals. Predicted values are controlled for all other covariates in the model (e.g., body mass, sampling time, etc.; set to their reference values, see material and methods). Colours indicate sex: females in red (and on the left), males in blue (on the right).

**Table 2:** Main model summary of model testing the effect of various traits on baseline levels of CORT. For the alternative model without the interaction of sex and breeding latitude see Supporting Information, Table S8. Values with CrI95 not containing zero are highlighted in **bold** and regarded as significant support for an effect. Weakly supported trends in models are highlighted in *italic*.

| predictor | baseline corticosterone |  |  |
| --- | --- | --- | --- |
|  | b | LCrI | UCrI |
| *ln species specific body mass* | *-0.19* | *-0.43* | *0.06* |
| IBMD | -0.02 | -0.06 | 0.03 |
| **blood sampling latency (min)** | **0.71** | **0.54** | **0.89** |
| **time since sunrise (h)** | **-0.05** | **-0.09** | **-0.01** |
| breeding status (nonbreeding) | 0.25 | -0.10 | 0.61 |
| **breeding latitude (tropical)** | **-0.88** | **-1.44** | **-0.34** |
| sex (female) | -0.20 | -0.53 | 0.13 |
| migration distance (1000km) | -0.03 | -0.18 | 0.13 |
| clutch size | -0.05 | -0.19 | 0.10 |
| **diet (carnivore)** | **0.12** | **0.03** | **0.21** |
| diet (nectarivore) | 0.06 | -0.05 | 0.18 |
| *breeding latitude (tropical) × sex (female)* | *0.40* | *-0.03* | *0.83* |
| *effect of phylogeny* | *0.10* | *0.00* | *0.33* |
| *Bayesian conditional R<sup>2</sup>* | *0.36* | *0.27* | *0.44* |
| *Bayesian marginal R<sup>2</sup>* | *0.31* | *0.23* | *0.38* |

**Table 3:**
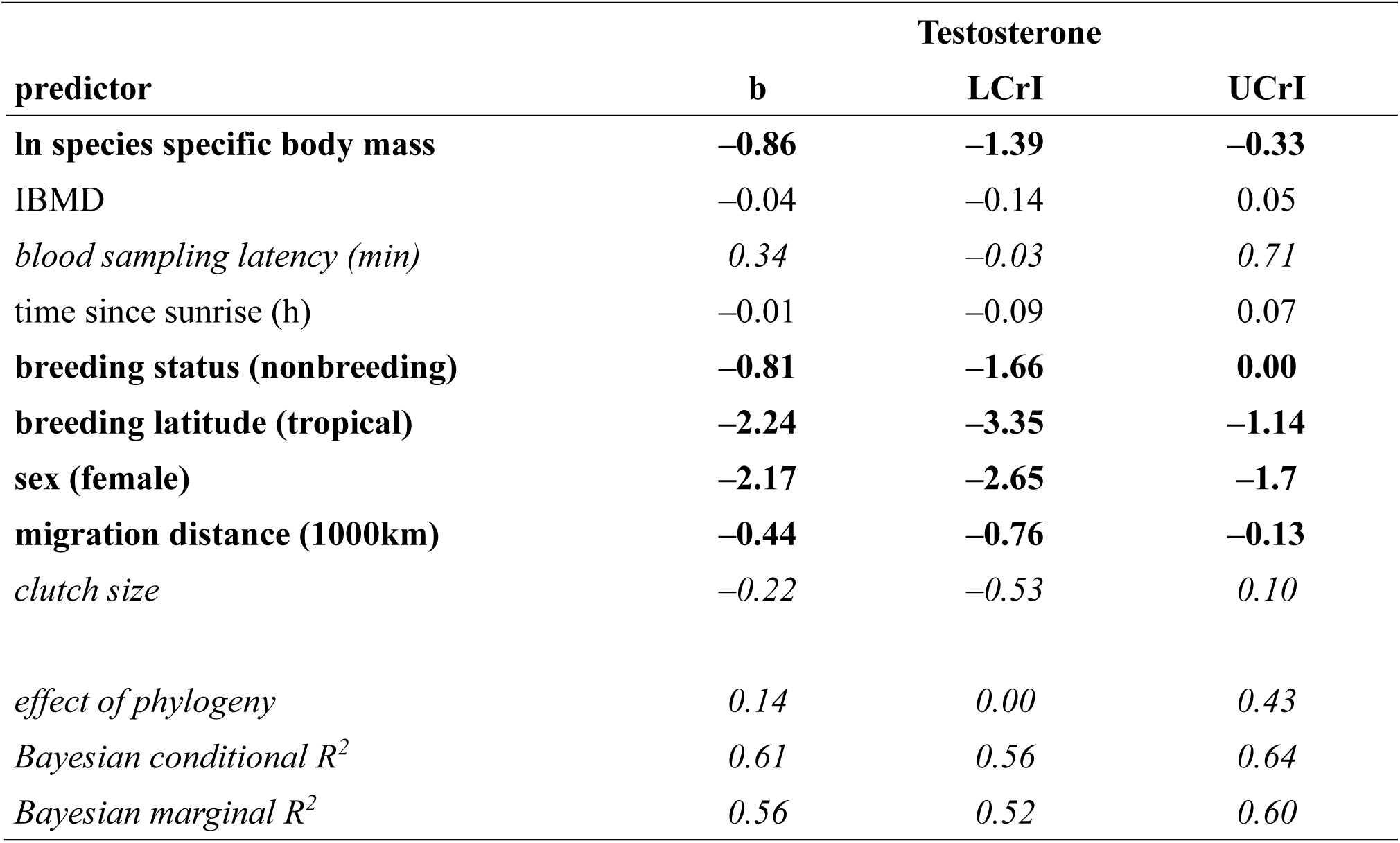
Summary of main model testing the effect of various traits on TEST. For the alternative model with the interaction of sex and breeding latitude see Supporting Information, Table S13. Values with CrI95 not containing zero are highlighted in **bold** and regarded as significant support for an effect. Weakly supported trends in models are highlighted in *italic*.

| predictor | b | Testosterone |  |
| --- | --- | --- | --- |
|  |  | LCrI | UCrI |
| **ln species specific body mass** | **−0.86** | **−1.39** | **−0.33** |
| IBMD | −0.04 | −0.14 | 0.05 |
| *blood sampling latency (min)* | *0.34* | *−0.03* | *0.71* |
| time since sunrise (h) | −0.01 | −0.09 | 0.07 |
| **breeding status (nonbreeding)** | **−0.81** | **−1.66** | **0.00** |
| **breeding latitude (tropical)** | **−2.24** | **−3.35** | **−1.14** |
| **sex (female)** | **−2.17** | **−2.65** | **−1.7** |
| **migration distance (1000km)** | **−0.44** | **−0.76** | **−0.13** |
| *clutch size* | *−0.22* | *−0.53* | *0.10* |
| *effect of phylogeny* | *0.14* | *0.00* | *0.43* |
| *Bayesian conditional R<sup>2</sup>* | *0.61* | *0.56* | *0.64* |
| *Bayesian marginal R<sup>2</sup>* | *0.56* | *0.52* | *0.60* |

While a positive correlation between CORT and clutch size was detected in an ASE model (Supporting Information, Table S7), this association did not persist in the main model. Baseline CORT was associated neither with mean species body mass nor the individual deviation from this mean in any of the models, although there was a trend for lower levels in larger species. Across species, baseline CORT levels increased with the proportion of carnivorous diet but not with the nectarivorous/frugivorous diet, meaning species with more animal matter in their diet had slightly higher baseline CORT. Finally, there was no association with migration distance in the model. To explore migration in greater detail, we analysed the association between CORT and migration only in temperate species, but again found no significant relationship (Supporting information, Table S9)

In a subset of tropical species (55 species), we evaluated the effect of elevation. The model showed no association with either the species-specific elevational midpoint or the difference between capture elevation and this midpoint. In this model, baseline CORT was higher in females than males and was higher during the nonbreeding season than during the breeding season (see Supporting Information, Table S10).

The stress-induced CORT was positively correlated with baseline CORT levels. The stress-induced CORT reactivity (i.e., stress-induced CORT level controlled for the baseline level) was negatively associated with migration distance (controlled for breeding latitude), did not differ between breeding latitudes, and was not associated with sex (Supporting Information, Table S11). In a model with baseline blood glucose levels as a dependent variable, this trait was not associated with baseline CORT levels (Supporting Information, Table S12).

### b) Testosterone

Similarly to CORT, we first tested whether breeding latitude influenced sexual differences in TEST by comparing models with and without a sex × breeding latitude interaction. The interaction was not statistically supported (95% CrI: –0.72; 1.11), and the model including it showed a slightly worse fit than the simpler model (*Δelpd loo* = –1.3, s.d. = 0.5; Table 3). We therefore used the simpler model as our main model, with results summarised in Table 3, results of the alternative model in Supporting Information, Table S13. Results of ASE models are provided in Supporting Information, Table S7.

In the main model, breeding latitude (Cameroon versus Czechia) emerged as an important predictor, with tropical species showing lower TEST concentrations than temperate zone ones, and males consistently having higher levels than females (Fig. 2b). TEST levels were negatively associated with species specific body mass, but not with the IBMD. Clutch size was positively associated with TEST in the ASE model (Supporting Information, Table S7) but it lost its significance when breeding latitude and all other covariates were included in the main model. A negative correlation between TEST and migration distance was detected in both the full dataset (Table 3) and in a subset restricted to temperate species only (Table 4, Fig. 3).

**Figure 3:**
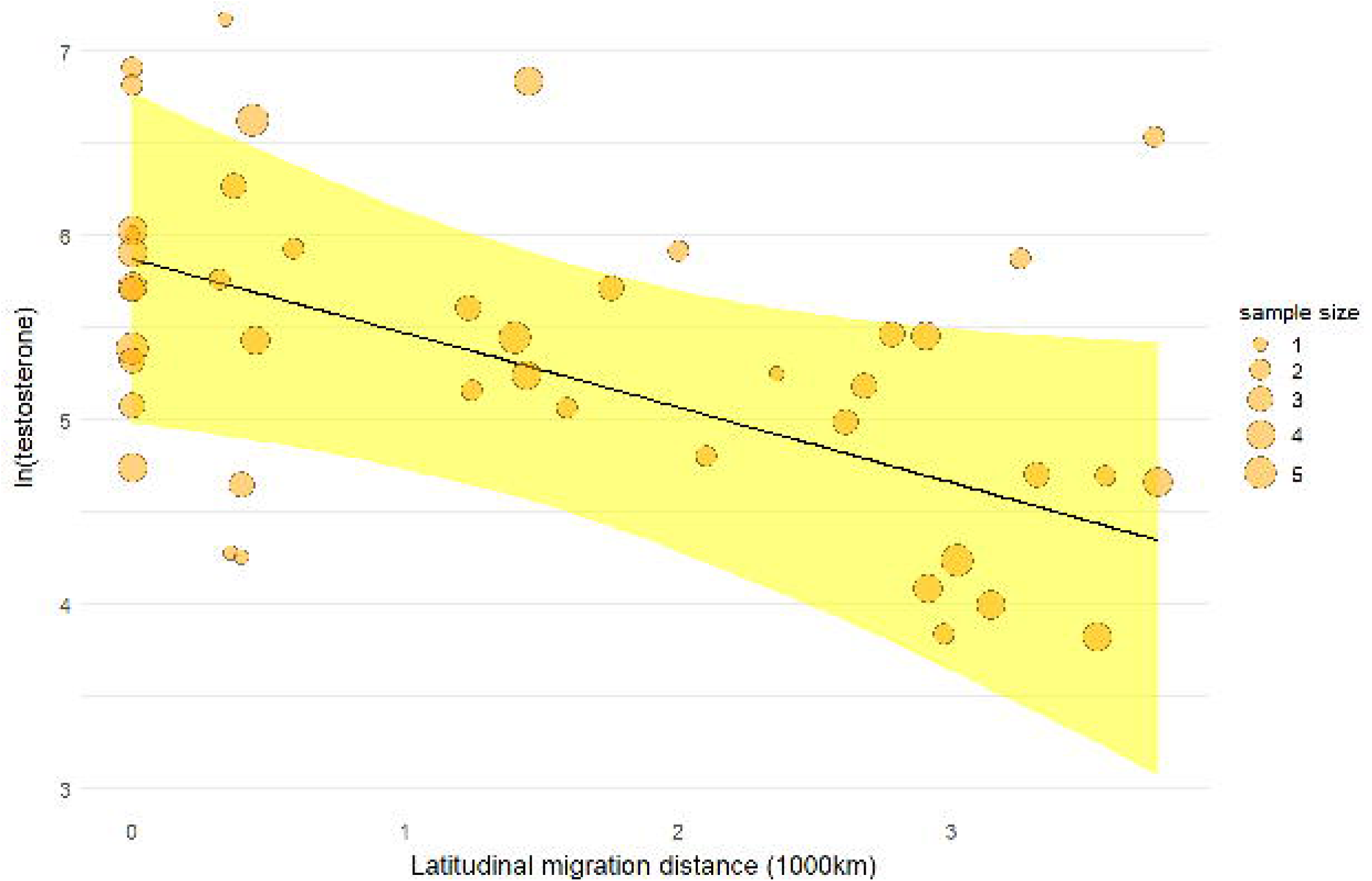
Relationship between migration distance (difference between the breeding and wintering latitudes) and plasma TEST for temperate species (for details of the model see Table 4). Circles represent TEST averages per species with size representing sample size.

**Table 4:** Model summary testing the association between TEST, sex and migration limited to temperate species only (n = 130 individuals of 44 species). Values with CrI95 not containing zero are highlighted in **bold** and regarded as significant support for an effect.

| predictor | b | Testosterone |  |
| --- | --- | --- | --- |
|  |  | LCrI | UCrI |
| ln species specific body mass | −0.40 | −1.30 | 0.49 |
| IBMD | −0.01 | −0.15 | 0.14 |
| time since sunrise (h) | 0.06 | −0.08 | 0.21 |
| blood sampling latency (min) | 0.49 | −0.12 | 1.12 |
| **sex (female)** | **−2.35** | **−3.19** | **−1.57** |
| **migration distance (1000km)** | **−0.41** | **−0.85** | **−0.01** |
| *effect of phylogeny* | *0.18* | *0.00* | *0.53* |
| *Bayesian conditional R<sup>2</sup>* | *0.54* | *0.43* | *0.62* |
| *Bayesian marginal R<sup>2</sup>* | *0.46* | *0.34* | *0.54* |

In a subset of 55 tropical species, TEST was higher in tropical males than females, but, as in CORT, showed no significant relationship with either species specific elevation midpoint or the difference between the elevation of capture and the midpoint. The model, however, suggested a trend of elevated plasma TEST during the breeding season in tropical species (Supporting Information, Table S10).

## Discussion

In this study, we present results of a broad-scale comparison of circulating plasma testosterone (TEST), baseline corticosterone (CORT), and stress-induced CORT reactivity, while controlling for key ecological and life history variables. All three measurements showed levels of repeatability comparable to those reported elsewhere for steroid hormones (Romero & Reed, 2008; Hau & Goymann, 2015; Taff *et al*., 2018) and for other physiological variables such as glucose (Tomášek *et al*., 2022). To the best of our knowledge, this is one of the first field studies (i.e., those analysing data collected according to a standardised protocol, rather than combining data from other studies) to examine interspecific variation in both steroid hormones across a broad range of avian families in temperate zone and the tropics, and the first to do so outside the Americas. Our results reveal consistent latitudinal differences: tropical species exhibited lower levels of CORT and TEST, corroborating previous findings from the Nearctic-Neotropic system (Hau *et al*., 2010) and other studies based on literature data (e.g. Goymann *et al*., 2004; Casagrande *et al*., 2018; Husak *et al*., 2021). Variation in TEST concentrations was further explained by body mass, migration distance, and sex, and variations in CORT also by diet. However, neither hormone was associated with individual deviation from species-specific body mass, possibly because sample size per species was insufficient to detect such an association. By extending the research to the understudied Afrotropical-Palearctic system, our study provides a critical test to the generality of previous evidence for latitudinal changes in steroid hormone concentrations in birds.

Lower levels of circulating CORT in Afrotropical than European temperate species may reflect the generally lower metabolic rates of tropical species compared to those from the temperate zone (Wiersma *et al*., 2007a, b; Bushuev *et al*., 2018), which would align with the primarily metabolic functions of CORT at baseline levels (Sapolsky *et al*., 2000). Although previous studies have reported a negative correlation between CORT and body size (Bókony *et al*., 2009; Hau *et al*., 2010; Vitousek *et al*., 2019), this relationship was only a weak trend in our main model. This implies that the relationship between CORT and body mass might not be universal. Similar to findings across all vertebrates (Vitousek *et al*., 2019; but see Casagrande *et al*., 2018), we found no effect of sex on baseline CORT in the main model. However, in tropical species, males tended to have lower CORT than females (Table 2, Fig. 2a, Supporting Information, Table S10). A similar pattern, i.e., higher levels in females than in males in tropical species, has previously been reported for baseline glucose (Tomášek *et al*., 2022). However, our data do not reveal a direct association between CORT and baseline glucose, despite baseline CORT being considered a metabolic hormone (Sapolsky *et al*., 2000; Jimeno & Verhulst, 2023). This may reflect the complex, context-dependent role of glucocorticoids, which can vary across ecological or physiological settings and among species that differ in their sensitivity to CORT, for example through different number of CORT receptors in their tissues (Sapolsky *et al*., 2000).

The difference in CORT between latitudes was greater in males than in females, although this latitude-sex interaction was only weakly supported. Low CORT in tropical males may reflect their prolonged and less synchronized breeding seasons and generally low population densities, which have been hypothesised to reduce metabolic demands due to fewer agonistic interactions between neighbouring breeding pairs (Stutchbury & Morton, 2022). By contrast, the more similar CORT in tropical and temperate zone females might reflect general female-specific energy demanding reproductive adaptations, such as egg production (Stutchbury & Morton, 1995). Surprisingly, however, clutch size itself, a widely used proxy for female reproductive investments, was not associated with CORT levels after accounting for differences in clutch sizes between tropical and temperate regions. This is unexpected, as a previous study have found a positive relationship between baseline CORT and brood value (composite of clutch size and species lifespan in Bókony *et al*., 2009). On the other hand, a study by Casagrande *et al*. (2018) reported a positive relationship between clutch size and the magnitude of change of baseline CORT between breeding and non-breeding season in passerines, rather than with breeding CORT alone. This might better reflect overall metabolic investment in reproduction. Unfortunately, we lack the non-breeding values of baseline CORT for temperate zone species and for part of the tropical ones to properly test this assumption. Still, baseline CORT was higher in the non-breeding season in tropical species, paralleling increases in baseline blood glucose (Tomášek *et al*., 2022) and potentially reflecting the energetic demands of moult (Lindström *et al*., 1993) rather than reproduction. Taken together, unlike in glucose (Tomášek *et al*., 2022), clutch size does not seem to be a primary driver in the difference in baseline CORT between tropical and temperate species.

Unlike in our previous study of blood glucose, there was no association of CORT levels with frugivory and nectarivory. This might suggest that high sugar availability in nectar can reduce reliance on gluconeogenesis in frugivorous and nectarivorous passerines, in a similar manner as in hummingbirds (McWhorter & Martínez Del Rio, 1999). On the other hand, species with a higher proportion of animal diet (mostly invertebrates in our dataset), low in simple sugars, had higher CORT levels. This likely reflects both the role of CORT in increased gluconeogenesis and the greater energetic unpredictability of carnivorous foraging (McEwen & Wingfield, 2003; Landys *et al*., 2006; Szarka & Lendvai, 2024). Together, these results highlight how dietary niche can shape endocrine traits via metabolic and ecological mechanisms.

CORT plays a key role in preparing birds for migration (Uehling *et al*., 2024), an energetically demanding life history stage that often immediately follows or precedes the breeding season. Within species, migratory populations may exhibit elevated CORT levels during the breeding season compared to sedentary populations (Krause *et al*., 2021; Gonzalez-Gomez *et al*., 2023). However, across temperate zone species, we found no association between breeding season baseline CORT and migration distance, consistent with previous analysis involving literature data sampled around the globe (Uehling *et al*., 2024). This may reflect flexible regulation of CORT (i.e., “CORT phenotypes”) across life history stages (Romero, 2002). Interestingly, and in contrast to Uehling *et al*. (2024), we found a negative association between migration distance and stress-induced CORT reactivity, suggesting a potential constraint on acute stress responsiveness in long-distance migrants, a pattern that merits further investigation. Stress-induced CORT reactivity and baseline CORT showed a positive relationship, consistent with Vitousek *et al*. (2019). No other ecological or life history traits were correlated with stress-induced CORT reactivity. However, compared to baseline levels dataset, our stress-induced CORT reactivity data included fewer species and individuals (75 individuals of 24 species), which may have reduced our ability to detect broader patterns.

Regarding testosterone, tropical birds exhibited lower circulating TEST concentrations than temperate ones, and TEST scaled negatively with species body mass. This pattern is consistent with results from a field study of New World passerines (Hau *et al*., 2010) as well as studies compiling data from other sources (Garamszegi *et al*., 2005, 2008; Husak *et al*., 2021; although Afrotropical species were largely omitted from those datasets; table 3). Such relationship may reflect the negative allometry between body mass and testes size, the primary organs responsible for TEST production (MacLeod & MacLeod, 2009).

As expected, TEST levels were higher in males, in line with its established role in regulating male breeding-related phenotypes (Wingfield *et al*., 2001), Fig. 2b). Sex difference in TEST levels was maintained and of a similar magnitude even among tropical species, where females often participate in territorial defence and territorial singing (Mikula *et al*., 2019), which is expected to select for more similar TEST levels across sexes (Møller *et al*., 2005). It should be noted that our temperate zone data include only individuals sampled during the breeding season, whereas tropical samples included individuals from both breeding and non-breeding seasons, although this has been controlled for in the main model. Even though many tropical species exhibit prolonged and opportunistic breeding (Stutchbury & Morton, 2022), distinct seasonal peaks in reproductive activity are still common (Goymann *et al*., 2006; Chiver *et al*., 2014; Lopes *et al*., 2021) and our results also indicate a tendency for elevated TEST among males of tropical species sampled during the breeding season (Supporting Information, Table S10). Neglecting breeding status may therefore bias comparative analyses of hormone levels across latitudes.

To the best of our knowledge, our study is the first to directly test the association between TEST and clutch size, a commonly used proxy for reproductive investment in birds (see above). In an ASE model (Supporting Information, Table S7), clutch size was indeed positively correlated with TEST. However, in our main model controlling for breeding latitude, this relationship disappeared, likely due to the systematically smaller clutch sizes of tropical species compared to temperate ones (Jetz *et al*., 2008; table 3). Hence, as with CORT, clutch size does not appear to be a primary driver explaining interspecific variation in latitudinal differences in TEST.

Our results, both from ASE and multivariate models, reveal a clear negative association between TEST and migration distance (Tables 3 and 4, Fig. 3). This is somewhat unexpected, as migratory species typically have shorter breeding seasons than sedentary ones, and species with shorter breeding seasons, as typically inferred from tropical and temperate species comparisons, are expected to have higher breeding TEST levels (Hau *et al*., 2010; Husak *et al*., 2021). However, migratory passerines in the northern temperate regions tend to have smaller clutches (Bruderer & Salewski, 2009; but see above regarding the association between clutch size and TEST in the whole dataset), higher interannual survival than their sedentary counterparts (Winger & Pegan, 2021) and lower basal metabolic rates (Wiersma *et al*., 2007b). In these life-history traits, long-distance migratory species thus seem to resemble tropical species which may explain their lower TEST (but see Soriano-Redondo *et al*., 2020). Although a higher TEST was reported for migratory than sedentary species previously (Garamszegi *et al*., 2008), unlike that study we quantified migration distance as a continuous variable, providing a finer description of variation in migration strategies than a coarse division into migrants and sedentary species. This approach allows a better reflection of the possibility that short distance migrants resemble sedentary species in many life history traits rather than long-distance migrants (Horák *et al*., 2022), but given the contradictory results, further research in this field is needed to clarify this relationship.

Considering tropical species only, we did not find any evidence for change in TEST with elevation, consistent with the pattern observed in CORT, and contrary to the positive trend observed for TEST in Neotropics (Goymann *et al*., 2004). Compared to the aforementioned study, which used data from studies ranging from 0 to ca. 3500 m a.s.l., our elevational range is limited by the tree line on Mt.

Cameroon which is located at 2300 m a.s.l. (Pernice *et al*., 2025). Nevertheless, these findings indicate more research is needed, as our previous study, using a slightly expanded number of species from the same locality, detected a positive correlation between blood glucose and altitude (Tomášek *et al*., 2022), even on this shorter elevational gradient.

## Conclusion

Despite their frequent phylogenetic proximity to well-studied European temperate taxa, Afrotropical passerine species remain underrepresented in comparative research. This study provides the first standardized sampling of circulating steroid hormones across Afrotropical and European temperate passerines, and one of the few such datasets worldwide. Our findings indicate that reduced levels of CORT and TEST represent a general phenomenon, that has evolved independently in tropical regions across continents and phylogenetic lineages. This outcome may not be surprising, as both hormones are expected to co-evolve with other life history traits associated with the pace of life, being reduced in species with slower life histories. Although direct data on basal metabolic rates are not yet available for Afrotropical taxa, indirect evidence (e.g. Jetz *et al*., 2008; Albrecht *et al*. 2013; Stevens *et al*., 2013; Valcu *et al*., 2014; Tomášek *et al*., 2022) do indeed suggest that they, like their Neotropical (Wiersma *et al*., 2007b) and Southeast Asian (Bushuev *et al*., 2018) counterparts, exhibit a relatively slow pace of life.

Our study is also the first to account for both intra- and interspecific variability in steroid hormone levels. In particular, our comparison of hormone levels in sexes within the tropical-temperate framework provides novel insights into the evolution of hormonal profiles in females regarding to life history traits and underscores the importance of including both sexes to obtain unbiased estimates as well as control for potentially confounding variables like season, sampling time or time of day. Furthermore, our results based on a fine scoring of migration challenges previous report that migratory species have lower TEST than sedentary ones. Future research should quantify additional physiological traits in Afrotropical taxa, particularly basal metabolic rates, and extend sampling to underexplored regions, such as the southern temperate zone, where life history pace and metabolic rates may diverge (Bech *et al*., 2016).

## Data Availability Statement

All data and script used to generate the results were uploaded to figshare and can be accessed here: https://doi.org/10.6084/m9.figshare.30073582

## Acknowledgements

We would like to thank to everyone who helped with the data collection in the field, namely Jaromír Čejka, Marek Brindzák and Kateřina Šimonová. Also, we are very grateful to the Chief of Bokwaongo, Chief of Bakingili and Bokwaongo community for their help with the field work in Cameroon.

## Author contributions

OK and TA conceived and designed the study, all authors performed research, OK analysed the data, OK and TA drafted the manuscript with input from OT, and all authors contributed to revisions of the manuscript.

## Ethics

The study was conducted in accordance with the Guidelines for Animal Care and Treatment of the European Union and was approved by the Animal Welfare Committee of the Czech Academy of Sciences under protocol number 09/2015 and the Cameroon Ministry of Research and Innovation under the research permit numbers 0021/MINRESI/B00/C00/C10/C14 and 0412/PRS/MINFOF/SG/DFAP/SDVEF/SC.

## References

1. Albrecht T, Kleven O, Kreisinger J, Laskemoen T, Omotoriogun TC, Ottosson U, Reif J, Sedláček O, Hořák D, Robertson RJ, Lifjeld JT. 2013. Sperm competition in tropical versus temperate zone birds. Proceedings of the Royal Society B: Biological Sciences 280: 20122434–20122434. 10.1098/rspb.2012.2434

2. Apfelbeck B, Helm B, Illera JC, Mortega KG, Smiddy P, Evans NP. 2017. Baseline and stress-induced levels of corticosterone in male and female Afrotropical and European temperate stonechats during breeding. BMC Evolutionary Biology 17: 114. 10.1186/s12862-017-0960-9

3. Bantock TM, Prys-Jones RP, Lee PLM. 2008. New and improved molecular sexing methods for museum bird specimens. Molecular Ecology Resources 8: 519–528. 10.1111/j.1471-8286.2007.01999.x

4. Barber RA, Yang J, Yang C, Barker O, Janicke T, Tobias JA. 2024. Climate and ecology predict latitudinal trends in sexual selection inferred from avian mating systems. PLOS Biology 22: e3002856. 10.1371/journal.pbio.3002856

5. Bauer CM, Watts HE. 2021. Corticosterone’s roles in avian migration: Assessment of three hypotheses. Hormones and Behavior 135: 105033. 10.1016/j.yhbeh.2021.105033

6. Bech C, Chappell MA, Astheimer LB, Londoño GA, Buttemer WA. 2016. A ‘slow pace of life’ in Australian old-endemic passerine birds is not accompanied by low basal metabolic rates. Journal of Comparative Physiology B: Biochemical, Systemic, and Environmental Physiology 186: 503–512.

7. Bicudo JEPW, Buttemer WA, Chappell MA, Pearson JT, Bech C. 2010. Ecological and Environmental Physiology of Birds. Oxford University Press.

8. Bílková Z, Adámková M, Albrecht T, Šimek Z. 2019. Determination of testosterone and corticosterone in feathers using liquid chromatography-mass spectrometry. Journal of Chromatography A 1590: 96–103. 10.1016/j.chroma.2018.12.069

9. BirdLife International and Handbook of the Birds of the World. 2018. Bird Species Distribution Maps of the World. Version 2018.1.

10. Bókony V, Lendvai ÁZ, Liker A, Angelier F, Wingfield JC, Chastel O. 2009. Stress Response and the Value of Reproduction: Are Birds Prudent Parents? The American Naturalist 173: 589–598. 10.1086/597610

11. Boyce AJ, Freeman BG, Mitchell AE, Martin TE. 2015. Clutch size declines with elevation in tropical birds. The Auk 132: 424–432. 10.1642/AUK-14-150.1

12. Bruderer B, Salewski V. 2009. Lower annual fecundity in long-distance migrants than in less migratory birds of temperate Europe. Journal of Ornithology 150: 281–286. 10.1007/s10336-008-0348-0

13. Bürkner PC. 2017. brms: An *R* Package for Bayesian Multilevel Models Using *Stan*. Journal of Statistical Software 80. 10.18637/jss.v080.i01

14. Bushuev A, Tolstenkov O, Zubkova E, Solovyeva E, Kerimov A. 2018. Basal metabolic rate in free-living tropical birds: the influence of phylogenetic, behavioral, and ecological factors. Current Zoology 64: 33–43. 10.1093/cz/zox018

15. Canoine V, Fusani L, Schlinger B, Hau M. 2007. Low sex steroids, high steroid receptors: Increasing the sensitivity of the nonreproductive brain. Developmental Neurobiology 67: 57–67. 10.1002/dneu.20296

16. Casagrande S, Zsolt Garamszegi L, Goymann W, Donald J, Francis CD, Fuxjager MJ, Husak JF, Johnson MA, Kircher B, Knapp R, Martin LB, Miller ET, Schoenle LA, Vitousek MN, Williams TD, Hau M. 2018. Do Seasonal Glucocorticoid Changes Depend on Reproductive Investment? A Comparative Approach in Birds. Integrative and Comparative Biology 58. 10.1093/icb/icy022

17. Cepák J, Klvaňa P, Škopek J, Schröpfer L, Jelínek M, Hořák D, Formánek J, Zárybnický J. 2008. Atlas migrace ptáků České republiky a Slovenska. Prague: Aventinum.

18. Chiver I, Stutchbury BJM, Morton ES. 2014. Seasonal variation in male testosterone levels in a tropical bird with year-round territoriality: Testosterone Levels in Red-throated Ant-tanagers. Journal of Field Ornithology 85: 1–9. 10.1111/jofo.12044

19. Cooney CR, He Y, Varley ZK, Nouri LO, Moody CJA, Jardine MD, Liker A, Székely T, Thomas GH. 2022. Latitudinal gradients in avian colourfulness. Nature Ecology & Evolution 6: 622–629. 10.1038/s41559-022-01714-1

20. Dawson A. 2008. Control of the annual cycle in birds: endocrine constraints and plasticity in response to ecological variability. Philosophical Transactions of the Royal Society B: Biological Sciences 363: 1621–1633. 10.1098/rstb.2007.0004

21. Deviche P, Valle S, Gao S, Davies S, Bittner S, Carpentier E. 2016. The seasonal glucocorticoid response of male Rufous-winged Sparrows to acute stress correlates with changes in plasma uric acid, but neither glucose nor testosterone. General and Comparative Endocrinology 235: 78–88. 10.1016/j.ygcen.2016.06.011

22. Foo YZ, Nakagawa S, Rhodes G, Simmons LW. 2017. The effects of sex hormones on immune function: a meta-analysis. Biological Reviews 92: 551–571. 10.1111/brv.12243

23. Garamszegi L, Eens M, Hurtrezbousses S, Moller A. 2005. Testosterone, testes size, and mating success in birds: a comparative study. Hormones and Behavior 47: 389–409. 10.1016/j.yhbeh.2004.11.008

24. Garamszegi LZ, Hirschenhauser K, Bokony V, Eens M, Hurtrez-Bousses S, Moller AP, Oliveira RF, Wingfield JC. 2008. Latitudinal distribution, migration, and testosterone levels in birds. American Naturalist 172: 533–546. 10.1086/590955

25. Ghalambor CK, Martin TE. 2000. Parental investment strategies in two species of nuthatch vary with stage-specific predation risk and reproductive effort. Animal Behaviour 60: 263–267. 10.1006/anbe.2000.1472

26. Gonzalez-Gomez PL, Villavicencio CP, Quispe R, Schwabl P, Cornelius JM, Ramenofsky M, Krause JS, Wingfield JC. 2023. Perspectives on environmental heterogeneity and seasonal modulation of stress response in neotropical birds. Hormones and Behavior 152: 105359. 10.1016/j.yhbeh.2023.105359

27. Goymann W, Geue D, Schwabl I, Flinks H, Schmidl D, Schwabl H, Gwinner E. 2006. Testosterone and corticosterone during the breeding cycle of equatorial and European stonechats (*Saxicola torquata axillaris* and *S. t. rubicola*). Hormones and Behavior 50: 779–785. 10.1016/j.yhbeh.2006.07.002

28. Goymann W, Moore IT, Scheuerlein A, Hirschenhauser K, Grafen A, Wingfield JC. 2004. Testosterone in tropical birds: effects of environmental and social factors. The American naturalist 164: 327–334. 10.1086/422856

29. Griffiths R, Double MC, Orr K, Dawson RJG. 1998. A DNA test to sex most birds. Molecular Ecology 7: 1071–1075. 10.1046/j.1365-294x.1998.00389.x

30. Hau M, Goymann W. 2015. Endocrine mechanisms, behavioral phenotypes and plasticity: known relationships and open questions. Frontiers in Zoology 12: S7. 10.1186/1742-9994-12-S1-S7

31. Hau M, Ricklefs RE, Wikelski M, Lee KA, Brawn JD. 2010. Corticosterone, testosterone and life-history strategies of birds. Proceedings. Biological sciences / The Royal Society 277: 3203–12. 10.1098/rspb.2010.0673

32. Hille SM, Cooper CB. 2015. Elevational trends in life histories: revising the pace-of-life framework. Biological Reviews 90: 204–213. 10.1111/brv.12106

33. Horák K, Bobek L, Adámková M, Kauzál O, Kauzálová T, Manialeu JP, Nguelefack TB, Nana ED, Jønsson KA, Munclinger P, Hořák D, Sedláček O, Tomášek O, Albrecht T. 2022. Feather growth and quality across passerines is explained by breeding rather than moulting latitude. Proceedings of the Royal Society B: Biological Sciences 289. 10.1098/rspb.2021.2404 https://doi.org/10.1093/zoolinnean/zlaf123

34. Horák K, Kotasová-Adámková M, Tomášek O, Kauzál O, Kauzálová T, Mani FT, Chi JE, Albrecht T. in press. Afrotropical passerines grow wing feathers faster than their European counterparts. Zoological Journal of the Linnean Society. 10.1093/zoolinnean/zlaf123

35. Husak JF, Fuxjager MJ, Johnson MA, Vitousek MN, Donald JW, Francis CD, Goymann W, Hau M, Kircher BK, Knapp R, Martin LB, Miller ET, Schoenle LA, Williams TD. 2021. Life history and environment predict variation in testosterone across vertebrates. Evolution 75: 1003–1010. 10.1111/evo.14216

36. Jessop TS, Woodford R, Symonds MRE. 2013. Macrostress: do large-scale ecological patterns exist in the glucocorticoid stress response of vertebrates? Functional Ecology 27: 120–130. 10.1111/j.1365-2435.2012.02057.x

37. Jetz W, Sekercioglu CH, Böhning-Gaese K. 2008. The Worldwide Variation in Avian Clutch Size across Species and Space. PLoS Biology 6: e303. 10.1371/journal.pbio.0060303

38. Jimenez AG, Harper JM, Queenborough SA, Williams JB. 2012. Linkages between the life-history evolution of tropical and temperate birds and the resistance of their cells to oxidative and non-oxidative chemical injury. Journal of Experimental Biology: jeb.079889. 10.1242/jeb.079889

39. Jimeno B, Verhulst S. 2023. Meta-analysis reveals glucocorticoid levels reflect variation in metabolic rate, not ‘stress’. eLife 12: RP88205. 10.7554/eLife.88205

40. Krause JS, Németh Z, Pérez JH, Chmura HE, Word KR, Lau HJ, Swanson RE, Cheah JC, Quach LN, Meddle SL, Wingfield JC, Ramenofsky M. 2021. Annual regulation of adrenocortical function in migrant and resident subspecies of white-crowned sparrow. Hormones and Behavior 127: 104884. 10.1016/j.yhbeh.2020.104884

41. Landys MM, Ramenofsky M, Wingfield JC. 2006. Actions of glucocorticoids at a seasonal baseline as compared to stress-related levels in the regulation of periodic life processes. General and Comparative Endocrinology 148: 132–149. 10.1016/j.ygcen.2006.02.013

42. Lindström Å, Visser GH, Daan S. 1993. The Energetic Cost of Feather Synthesis Is Proportional to Basal Metabolic Rate. Physiological Zoology 66: 490–510. 10.1086/physzool.66.4.30163805

43. Lisovski S, Neumann R, Albrecht T, Munclinger P, Ahola MP, Bauer S, Cepak J, Fransson T, Jakobsson S, Jaakkonen T, Klvana P, Kullberg C, Laaksonen T, Metzger B, Piha M, Shurulinkov P, Stach R, Ström K, Velmala W, Briedis M. 2021. The Indo-European flyway: Opportunities and constraints reflected by Common Rosefinches breeding across Europe. Journal of Biogeography 48: 1255–1266. 10.1111/jbi.14085

44. Lopes LE, Teixeira JPG, Meireles RC, Bastos DSS, De Oliveira LL, Solar R. 2021. High Seasonal Variation of Plasma Testosterone Levels for a Tropical Grassland Bird Resembles Patterns of Temperate Birds. Physiological and Biochemical Zoology 94: 143–151. 10.1086/713503

45. Lüdecke D, Ben-Shachar M, Patil I, Waggoner P, Makowski D. 2021. performance: An R Package for Assessment, Comparison and Testing of Statistical Models. Journal of Open Source Software 6: 3139. 10.21105/joss.03139

46. MacLeod CD, MacLeod RC. 2009. The relationship between body mass and relative investment in testes mass in amniotes and other vertebrates. Oikos 118: 903–916. 10.1111/j.1600-0706.2008.17426.x

47. Maness TJ, Grace JK, Hirchak MR, Tompkins EM, Anderson DJ. 2023. Circulating corticosterone predicts near-term, while H/L ratio predicts long-term, survival in a long-lived seabird. Frontiers in Ecology and Evolution 11. 10.3389/fevo.2023.1172904

48. Martin TE, Riordan MM, Repin R, Mouton JC, Blake WM. 2017. Apparent annual survival estimates of tropical songbirds better reflect life history variation when based on intensive field methods. Global Ecology and Biogeography 26: 1386–1397. 10.1111/geb.12661

49. McEwen BS, Wingfield JC. 2003. The concept of allostasis in biology and biomedicine. Hormones and Behavior 43: 2–15. 10.1016/S0018-506X(02)00024-7

50. McWhorter TJ, Martínez Del Rio C. 1999. Food ingestion and water turnover in hummingbirds: how much dietary water is absorbed? Journal of Experimental Biology 202: 2851–2858. 10.1242/jeb.202.20.2851

51. Mikula P, Tószögyová A, Hořák D, Petrusková T, Storch D, Albrecht T. 2019. Female solo song and duetting are associated with different territoriality in songbirds. Behavioral Ecology 31: 322–329. 10.1093/beheco/arz193

52. Millanes PM, Pérez-Rodríguez L, Rubalcaba JG, Gil D, Jimeno B. 2024. Corticosterone and glucose are correlated and show similar response patterns to temperature and stress in a free-living bird. Journal of Experimental Biology 227: jeb246905. 10.1242/jeb.246905

53. Møller AP, Garamszegi LZ, Gil D, Hurtrez-Boussès S, Eens M. 2005. Correlated evolution of male and female testosterone profiles in birds and its consequences. Behavioral Ecology and Sociobiology 58: 534–544. 10.1007/s00265-005-0962-2

54. Moore IT, Vernasco BJ, Escallón C, Small TW, Ryder TB, Horton BM. 2019. Tales of testosterone: Advancing our understanding of environmental endocrinology through studies of neotropical birds. General and Comparative Endocrinology 273: 184–191. 10.1016/j.ygcen.2018.07.003

55. Muñoz AP, Kéry M, Martins PV, Ferraz G. 2018. Age effects on survival of Amazon forest birds and the latitudinal gradient in bird survival. The Auk 135: 299–313. 10.1642/AUK-17-91.1

56. Myhrvold NP, Baldridge E, Chan B, Sivam D, Freeman DL, Ernest SKM. 2015. An amniote life-history database to perform comparative analyses with birds, mammals, and reptiles. Ecology 96: 3109–3109. 10.1890/15-0846R.1

57. Pernice R, Sedláček O, Albrecht T, Tomášek O, Kauzál O, Kauzálová T, Motombi FN, Ewome FL, Ferenc M, Chmel K, Mlíkovský J, Riegert J, Kamga SM, Hořák D. 2025. Functional and Phylogenetic Structure of Forest Bird Assemblages Along an Afrotropical Elevational Gradient. Ecology and Evolution 15: e72065. 10.1002/ece3.72065

58. QGIS.org. 2025. QGIS Geographic Information System. Open Source Geospatial Foundation Project. https://qgis.org

59. R Core Team. 2023. R: A Language and Environment for Statistical Computing. https://www.r-project.org

60. Ricklefs RE, Wikelski M. 2002. The physiology/life-history nexus. Trends in Ecology and Evolution 17: 462–468. 10.1016/S0169-5347(02)02578-8

61. Romero LM. 2002. Seasonal changes in plasma glucocorticoid concentrations in free-living vertebrates. General and Comparative Endocrinology 128: 1–24. 10.1016/S0016-6480(02)00064-3

62. Romero LM, Reed JM. 2005. Collecting baseline corticosterone samples in the field: is under 3 min good enough? Comparative Biochemistry and Physiology Part A: Molecular & Integrative Physiology 140: 73–79. 10.1016/j.cbpb.2004.11.004

63. Romero LM, Reed JM. 2008. Repeatability of baseline corticosterone concentrations. General and Comparative Endocrinology 156: 27–33. 10.1016/j.ygcen.2007.10.001

64. Sapolsky RM, Romero LM, Munck AU. 2000. How Do Glucocorticoids Influence Stress Responses? Integrating Permissive, Suppressive, Stimulatory, and Preparative Actions*. Endocrine Reviews 21: 55–89. 10.1210/edrv.21.1.0389

65. Schemske DW, Mittelbach GG, Cornell HV, Sobel JM, Roy K. 2009. Is There a Latitudinal Gradient in the Importance of Biotic Interactions? Annual Review of Ecology, Evolution, and Systematics 40: 245–269. 10.1146/annurev.ecolsys.39.110707.173430

66. Skutch AF. 1949. Do tropical birds rear as many young as they can nourish? Ibis 91: 430–455. 10.1111/j.1474-919X.1949.tb02293.x

67. Soriano-Redondo A, Gutiérrez JS, Hodgson D, Bearhop S. 2020. Migrant birds and mammals live faster than residents. Nature Communications 11: 1–8. 10.1038/s41467-020-19256-0

68. Spottiswoode C, Møller AP. 2004. Extrapair paternity, migration, and breeding synchrony in birds. Behavioral Ecology 15: 41–57. 10.1093/beheco/arg100

69. Stevens MC, Ottosson U, McGregor R, Brandt M, Cresswell W. 2013. Survival rates in West African savanna birds. Ostrich 84: 11–25. 10.2989/00306525.2013.772544

70. Stutchbury BJ, Morton ES. 1995. The Effect of Breeding Synchrony On Extra-Pair Mating Systems in Songbirds. Behaviour 132: 675–690. 10.1163/156853995X00081

71. Stutchbury BJ, Morton ES. 2022. Behavioral Ecology of Tropical Birds. Academic Press.

72. Sur S, Tiwari J, Malik S, Stevenson T. 2025. Endocrine and molecular regulation of seasonal avian immune function. Philosophical Transactions of the Royal Society B: Biological Sciences 380: 20230507. 10.1098/rstb.2023.0507

73. Synek P, Popelková A, Koubínová D, Šťastný K, Langrová I, Votýpka J, Munclinger P. 2016. Haemosporidian infections in the Tengmalm’s Owl (*Aegolius funereus*) and potential insect vectors of their transmission. Parasitology Research 115: 291–298. 10.1007/s00436-015-4745-z

74. Szarka EZ, Lendvai ÁZ. 2024. Trophic guilds differ in blood glucose concentrations: a phylogenetic comparative analysis in birds. Proceedings of the Royal Society B: Biological Sciences 291: 20232655. 10.1098/rspb.2023.2655

75. Taff CC, Schoenle LA, Vitousek MN. 2018. The repeatability of glucocorticoids: A review and meta-analysis. General and Comparative Endocrinology 260: 136–145. 10.1016/j.ygcen.2018.01.011

76. Taff CC, Wingfield JC, Vitousek MN. 2022. The relative speed of the glucocorticoid stress response varies independently of scope and is predicted by environmental variability and longevity across birds. Hormones and Behavior 144: 105226. 10.1016/j.yhbeh.2022.105226

77. Tobias JA, Sheard C, Seddon N, Meade A, Cotton AJ, Nakagawa S. 2016. Territoriality, Social Bonds, and the Evolution of Communal Signaling in Birds. Frontiers in Ecology and Evolution 4: 1–15. 10.3389/fevo.2016.00074

78. Tomášek O, Bobek L, Kauzálová T, Kauzál O, Adámková M, Horák K, Kumar SA, Manialeu JP, Munclinger P, Nana ED, Nguelefack TB, Sedláček O, Albrecht T. 2022. Latitudinal but not elevational variation in blood glucose level is linked to life history across passerine birds. Ecology Letters: 1–14. 10.1111/ele.14097

79. Uehling JJ, Regnier E, Vitousek MN. 2024. Does Migration Constrain Glucocorticoid Phenotypes? Testing Corticosterone Levels during Breeding in Migratory Versus Resident Birds. Integrative And Comparative Biology 64: 1826–1835. 10.1093/icb/icae110

80. Valcu M, Dale J, Griesser M, Nakagawa S, Kempenaers B. 2014. Global gradients of avian longevity support the classic evolutionary theory of ageing. Ecography 37: 930–938. 10.1111/ecog.00929

81. Vehtari A, Gelman A, Gabry J. 2017. Practical Bayesian model evaluation using leave-one-out cross-validation and WAIC. Statistics and Computing 27: 1413–1432. 10.1007/s11222-016-9696-4

82. Vehtari A, Gelman A, Simpson D, Carpenter B, Bürkner PC. 2021. Rank-Normalization, Folding, and Localization: An Improved R^ for Assessing Convergence of MCMC (with Discussion). Bayesian Analysis 16. 10.1214/20-BA1221

83. Vitousek MN, Johnson MA, Donald JW, Francis CD, Fuxjager MJ, Goymann W, Hau M, Husak JF, Kircher BK, Knapp R, Martin LB, Miller ET, Schoenle LA, Uehling JJ, Williams TD. 2018. HormoneBase, a population-level database of steroid hormone levels across vertebrates. Scientific Data 5: 180097. 10.1038/sdata.2018.97

84. Vitousek MN, Johnson MA, Downs CJ, Miller ET, Martin LB, Francis CD, Donald JW, Fuxjager MJ, Goymann W, Hau M, Husak JF, Kircher BK, Knapp R, Schoenle LA, Williams TD. 2019. Macroevolutionary Patterning in Glucocorticoids Suggests Different Selective Pressures Shape Baseline and Stress-Induced Levels. The American Naturalist 193: 866–880. 10.1086/703112

85. Wiersma P, Chappell M a, Williams JB. 2007a. Cold- and exercise-induced peak metabolic rates in tropical birds. Proceedings of the National Academy of Sciences of the United States of America 104: 20866–20871. 10.1073/pnas.0707683104

86. Wiersma P, Muñoz-Garcia A, Walker A, Williams JB. 2007b. Tropical birds have a slow pace of life. Proceedings of the National Academy of Sciences of the United States of America 104: 9340–5. 10.1073/pnas.0702212104

87. Wilman H, Belmaker J, Simpson J, De La Rosa C, Rivadeneira MM, Jetz W. 2014. EltonTraits 1.0: Species-level foraging attributes of the world’s birds and mammals. Ecology 95: 2027–2027. 10.1890/13-1917.1

88. Winger BM, Pegan TM. 2021. Migration distance is a fundamental axis of the slow-fast continuum of life history in boreal birds. Ornithology 138: ukab043.

89. Wingfield JC, Lynn SE, Soma KK. 2001. Avoiding the ‘Costs’ of Testosterone: Ecological Bases of Hormone-Behavior Interactions. Brain, Behavior and Evolution 57: 239–251. 10.1159/000047243

90. Wingfield JC, Maney DL, Breuner CW, Jacobs JD, Lynn S, Ramenofsky M, Richardson RD. 1998. Ecological Bases of Hormone—Behavior Interactions: The “Emergency Life History Stage”. American Zoologist 38: 191–206. 10.1093/icb/38.1.191

