## Supplemental information for "Comparative analysis of plasma steroid hormone levels reveals lower concentrations in Afrotropical than Palearctic temperate zone passerines"

Electronic Supplementary Material

for

### Supplementary Materials and methods

#### Detailed information about measuring concentrations of both hormones using LC-ESI-MS/MS

Corticosterone (CORT) and testosterone (TEST) concentrations in plasma samples were measured using liquid chromatography–electrospray ionization–tandem mass spectrometry (LC-ESI-MS/MS). The method is based on method developed by Bílková *et al.* (2019) which utilizes hormone derivatization prior to their analysis and was adapted for plasma samples as follows: plasma samples were prepared using solid-phase extraction (SPE) with Supelclean LC-18 Tubes (1 ml, 100 mg). The columns were conditioned with 1 ml of 100% methanol followed by 1 ml of 10% methanol in water. A 10 µl aliquot of plasma, spiked with 10 µl of internal standard solution (testosterone-D5 and corticosterone-D8, each at 6 ng/ml), was loaded onto the column. The columns were subsequently washed with 1 ml of 30% methanol in water, and the analytes were eluted with 400 µl of 100% methanol, collected directly into conical vials. The eluates were evaporated to dryness under a gentle stream of nitrogen. Subsequently, 50 µl of a derivatization reagent working solution (700 µl Amplifex™ Keto Reagent and 700 µl Amplifex™ Keto Reagent Diluent - QAO) was added to each micro-vial, and the samples were vortex-mixed. Derivatization was carried out for 120 minutes at 65 °C. After incubation, 10 µl of 70% methanol was added, followed by vortex mixing.

Chromatographic analysis was performed using LC-ESI-MS/MS (Agilent 1290 Infinity II + QQQ6495A, Agilent Technologies, Inc., Santa Clara, California, USA). Separation was achieved on an Acquity UPLC BEH C18 column (1.7 µm, 2.1 × 100 mm; Waters) equipped with a VanGuard pre-column (1.7 µm, 2.1 × 5 mm; Waters), maintained at 40 °C. The mobile phase consisted of solvent A (0.1% formic acid in water) and solvent B (acetonitrile), with the autosampler solvent set to 90% methanol. The flow rate was 300 µl/min and the injection volume was 10 µl. The gradient started at 82% A and 18% B, held for 5 minutes, and then linearly shifted to 67% A / 33% B at 8 minutes, 56.8% A / 43.2% B at 11 minutes, and 5% A / 95% B at 13 minutes. This composition was held until 20 minutes, followed by re-equilibration to the initial conditions (82% A / 18% B) at 20.1 minutes, maintained until 26 minutes. Analytes were quantified using multiple reaction monitoring (MRM) in positive electrospray ionization (ESI) mode. The specific precursor and product ion transitions, along with their corresponding collision energies, are listed in table below:

**MRM transitions and parameters used for quantification of target compounds.**

| Compound | Precursor Ion (m/z) | Product Ion (m/z) | Collision Energy (eV) | Polarity | Role |
|---|---|---|---|---|---|
| Testosterone-D5-QAO | 408.4 | 349.2 | 32 | Positive | Qualifier |
| Testosterone-D5-QAO | 408.4 | 167.9 | 52 | Positive | Quantifier |
| Testosterone-D5-QAO | 408.4 | 155.2 | 56 | Positive | Qualifier |
| Testosterone-QAO | 403.5 | 344.2 | 28 | Positive | Qualifier |
| Testosterone-QAO | 403.5 | 164.1 | 52 | Positive | Quantifier |
| Testosterone-QAO | 403.5 | 152.1 | 48 | Positive | Qualifier |
| Corticosterone-D8-QAO | 292.2 | 375.3 | 24 | Positive | Quantifier |
| Corticosterone-D8-QAO | 292.2 | 262.8 | 12 | Positive | Qualifier |
| Corticosterone-D8-QAO | 292.2 | 59.1 | 24 | Positive | Qualifier |
| Corticosterone-QAO | 288.4 | 369.2 | 20 | Positive | Quantifier |
| Corticosterone-QAO | 288.4 | 258.7 | 12 | Positive | Qualifier |

#### Testing for potential effect of time of day, sampling time and latency before freezing

Because of known diurnal variation in CORT (Schwabl *et al.* 2016) and rapid onset of its increase due to the stress of capture (Romero & Reed 2005), each time an individual bird was captured, time of day was noted as well as time of sampling latency (time between the moment of capture and blood sampling: *blood sampling latency*). Then, *time from sunrise* was calculated as the difference between time of capture and time of sunrise for that locality and date, which calculated using package suncalc v. 0.5.1 (Thieurmel & Elmarhraoui 2017) in R v. 4.4.2 (R Core Team 2023). Before centrifugation, samples were stored on ice or in a camping fridge and were deep frozen in liquid nitrogen right after centrifugation. Time of freezing of each sample was noted and the difference between deep freezing and time of capture was calculated (*deep freezing latency*).

We then tested the potential effect of any of these variables on both hormones. There was a positive correlation with *blood sampling latency* in both hormones and a negative correlation with *time from sunrise* for baseline corticosterone, but none with *deep freezing latency* (table S5). All subsequent analyses of both CORT and TEST were then controlled for *blood sampling latency* and *time for sunrise*.

#### Testing for potential effect of playback

In many ornithological field studies, playback of the species-specific song is use to either quantify or elicit territorial behaviour in a species (De Rosa *et al.* 2022). This can potentially influence plasma TEST levels of sampled individuals (Apfelbeck *et al.* 2017; but see Deviche *et al.* 2006, 2014). In our dataset, only a small subset of individuals was trapped using playback (35 individuals of mostly temperate species), nonetheless we investigated the role of playback on both TEST and CORT concentrations. To test the effect of playback, a separate analysis was made, which included only species with individuals both trapped with and without the use of conspecific playback (37 individuals of 11 species). There was no association between playback and neither of the hormones in the data (table S6), therefore we did not control for playback in further analyses.

#### Values below limit of quantification (LOQ)

Due to the limited volume of the samples which are possible to obtain from small passerines, 130 out of the 301 samples of circulating plasma TEST measured were under quantification limit (LOQ = 60 pg/ml). Their distribution was non-random in the dataset but rather followed expected biological distribution (table S3). We employed several ways how to treat these values.

1) We used the *cens* function from the brms package (LOQ set to 60) to impute the missing data. This approach allows the Bayesian model to treat values below LOQ as left-censored data and estimate them as part of the posterior distribution accounting for their uncertainty (Bürkner 2017).
2) We estimated the mean and standard deviation of Gaussian distribution fitted to the ln transformed data using *survreg* function from the package survival v. 3.7-0, which accounts for left censored data when estimating the distribution (Therneau 2025). We then generated random values from this distribution using *rtruncnorm* function from package truncnorm v. 1.0-9 (Mersman *et al.* 2023), excluding values above LOQ, and used these values in the models.
3) We replaced all values below LOQ with half the LOQ (30 pg/ml), a commonly used approached for handling the left-censored data (Shoari & Dubé 2018).
4) We excluded all values below LOQ from the dataset and performed the models using only remaining observations reducing the dataset to 171 individuals of 87 species.

Despite using different imputation strategies for values below LOQ, all four main models performed similarly with comparable parameter estimates (c.f. table 3 in main text and table S4).

### Supplementary figures

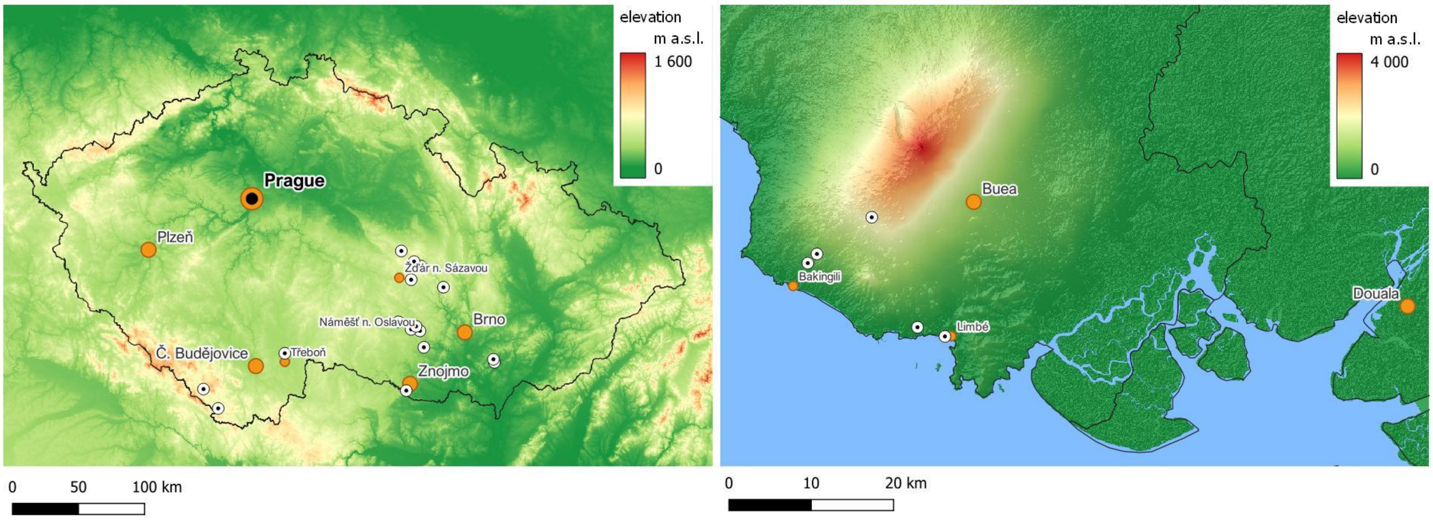

**Figure S1:** Approximate locations of sampling localities (dotted white circles) with position of major town and cities (orange circles). Left: Czechia (temperate zone), right: Cameroon (tropical zone).

### Supplementary tables

**Table S1:** List of fieldwork seasons and expeditions.

| site | year | months | note |
|---|---|---|---|
| Czechia (temperate) | 2018 | April – July | |
| | 2019 | March – June | |
| Cameroon (tropical) | 2017 | November – December | dry season |
| | 2018 | August – September<br>November – December | rainy season<br>dry season |

**Table S2:** List of species for which clutch size was not available in literature, the clutch size value used in the analysis instead and its source.

| species | clutch size | source |
|---|---|---|
| *Macrosphenus flavicans* | 1.9 | average of all species from the family Macrosphenidae from Gonzalez-Voyer *et al.* (2022) |
| *Urolais epichlorus* | 2 | average from two sister genera *Oreolais* and *Artisornis* from Billerman *et al.* (2022) |
| *Neocossyphus poensis* | 2 | clutch size of *Neocossyphus rufus* from Billerman *et al.* (2022) |
| *Saxicola torquatus* | 3.5 | Mackworth-Praed & Grant (1970) |
| *Deleornis fraseri* | 1.7 | average of all species of genus *Nectarinia s.l.* from Gonzalez-Voyer *et al.* (2022) |

**Table S3:** Number of TEST samples below the limit of quantification (LOQ = 60 pg/ml) out of the total number of samples for each group (breeding zone and sex). Percentages in parenthesis refer to the proportion of samples below LOQ for each group.

| sex | tropical | temperate | *SUBTOTAL* |
|---|---|---|---|
| male | 29/90 | 14/80 | *43/170 (25.3 %)* |
| female | 58/81 | 29/50 | *87/131 (66.4 %)* |
| *SUBTOTAL* | *87/171 (50.8 %)* | *43/130 (33.1 %)* | |

**Table S4:** Comparison of main model summaries of models testing the association between TEST and breeding latitude, sex, migration distance, clutch size and diet with different methods used to estimate values below LOQ: imputing random values based on observed gaussian distribution (see electronic supplementary material, Materials and methods), assigning the value of LOQ/2 = 30 pg/ml, and model using limited dataset without observations under LOQ (171 individuals of 87 species). Models were controlled for species specific body mass and the individual body mass deviation (IBMD), time from sunrise and blood sampling latency. Values with CrI95 not containing zero are highlighted in **bold** and regarded as significant support for an effect. Weakly supported trends in models are highlighted in *italic*. Underlined predictors were significant in the main model (main text, table 3).

All four models performed similarly. The most different model was the model using only values above the quantification limit with limited sample size. This is not surprising since the values below quantification limit were not randomly distributed but rather followed biologically meaningful expected distribution: more samples were absent in tropical rather than temperate species and more so in females than in males (table S3).

| **predictor** | **random** | | | **30 pg/ml** | | | **only values > LOQ** (n = 171) | | |
|---|---|---|---|---|---|---|---|---|---|
| | **b** | **LCrI** | **UCrI** | **b** | **LCrI** | **UCrI** | **b** | **LCrI** | **UCrI** |
| ln species specific body mass | **–0.8** | **–1.27** | **–0.32** | **–0.61** | **–0.97** | **–0.25** | **–0.44** | **–0.8** | **–0.09** |
| IBMD | –0.04 | –0.12 | 0.05 | –0.03 | –0.09 | 0.03 | –0.02 | –0.10 | 0.05 |
| blood sampling latency (min) | *0.36* | *–0.01* | *0.72* | **0.27** | **0.01** | **0.53** | 0.15 | –0.13 | 0.43 |
| time from sunrise (h) | 0.01 | –0.07 | 0.09 | 0.01 | –0.05 | 0.06 | 0.01 | –0.05 | 0.08 |
| breeding status (nonbreeding) | –0.39 | –1.08 | 0.30 | –0.30 | –0.82 | 0.21 | 0.20 | –0.49 | 0.89 |
| breeding latitude (tropical) | **–1.92** | **–2.93** | **–0.89** | **–1.72** | **–2.5** | **–0.95** | **–1.36** | **–2.12** | **–0.59** |
| sex (female) | **–1.69** | **–2.12** | **–1.25** | **–1.5** | **–1.82** | **–1.19** | **–1.09** | **–1.49** | **–0.69** |
| migration distance (1000km) | **–0.39** | **–0.69** | **–0.09** | **–0.32** | **–0.55** | **–0.1** | –0.08 | –0.30 | 0.14 |
| clutch size | –0.17 | –0.46 | 0.11 | *–0.15* | *–0.36* | *0.07* | –0.05 | –0.27 | 0.17 |

**Table S5:** Model summaries of models testing the effect of diurnal variation, and sampling and deep-freezing latency on baseline CORT and TEST. Values with CrI95 not containing zero are highlighted in **bold** and regarded as significant support for an effect.

| | Baseline corticosterone | | | Testosterone | | |
|---|---|---|---|---|---|---|
| **predictor** | **b** | **LCrI** | **UCrI** | **b** | **LCrI** | **UCrI** |
| time from sunrise (h) | **–0.05** | **–0.09** | **–0.01** | –0.07 | –0.18 | 0.04 |
| blood sampling latency (min) | **0.71** | **0.53** | **0.89** | **0.55** | **0.09** | **1.02** |
| deep freezing latency (min) | 0.08 | –0.10 | 0.26 | –0.39 | –0.91 | 0.11 |

**Table S6:** Model summary of the effect of using a playback on the levels of baseline CORT and TEST on a subset of species with individuals trapped both with the use of playback and without (n = 37 individuals of 11 species). Models were controlled for species specific body mass and the IBMD, and sex. Values with CrI95 not containing zero are highlighted in **bold** and regarded as significant support for an effect.

| | Baseline corticosterone | | | Testosterone | | |
|---|---|---|---|---|---|---|
| **predictor** | **b** | **LCrI** | **UCrI** | **b** | **LCrI** | **UCrI** |
| ln species specific body mass | 0.16 | –0.55 | 0.86 | –0.30 | –2.49 | 1.83 |
| IBMD | **0.13** | **0.02** | **0.23** | –0.13 | –0.46 | 0.15 |
| breeding latitude (tropical) | **–1.68** | **–3.24** | **–0.09** | –1.00 | –5.89 | 3.83 |
| sex (female) | –0.53 | –1.28 | 0.22 | **–2.70** | **–4.69** | **–0.92** |
| playback (yes) | –0.41 | –1.18 | 0.35 | 0.50 | –1.31 | 2.31 |

**Table S7:** Summary of predictor estimates of "adjusted single effect" (ASE) models for baseline CORT and TEST. All models were controlled for species specific body mass and IBMD, sex, blood sampling latency and time from sunrise. Values with CrI95 not containing zero are highlighted in **bold** and regarded as significant support for an effect. Weakly supported trends in models are highlighted in *italic*.

| | Baseline corticosterone | | | Testosterone | | |
|---|---|---|---|---|---|---|
| **predictor** | **b** | **LCrI** | **UCrI** | **b** | **LCrI** | **UCrI** |
| breeding latitude (tropical) | **–0.47** | **–0.74** | **–0.21** | **–1.27** | **–1.86** | **–0.69** |
| breeding status (nonbreeding) | 0.06 | –0.29 | 0.42 | **–1.39** | **–2.25** | **–0.57** |
| clutch size | **0.10** | **0.01** | **0.19** | **0.27** | **0.05** | **0.48** |
| diet (carnivore) | **0.08** | **0.01** | **0.15** | –0.12 | –0.30 | 0.06 |
| diet (herbivore) | **–0.09** | **–0.18** | **–0.02** | 0.14 | –0.06 | 0.34 |
| diet (nectarivore) | –0.02 | –0.11 | 0.08 | 0.14 | –0.08 | 0.37 |
| *temperate species only* | | | | | | |
| migration distance (1000km) | 0.10 | –0.06 | 0.25 | **–0.41** | **–0.84** | **–0.01** |
| clutch size | –0.06 | –0.25 | 0.13 | –0.39 | –0.88 | 0.10 |
| diet (carnivore) | **0.12** | **0.01** | **0.22** | –0.12 | –0.40 | 0.17 |
| diet (herbivore) | **–0.13** | **–0.23** | **–0.03** | 0.21 | –0.07 | 0.48 |
| diet (nectarivore) | 0.00 | –0.16 | 0.16 | –0.10 | –0.52 | 0.32 |
| *tropical species only* | | | | | | |
| species elevation midpoint | –0.05 | –0.32 | 0.24 | –0.22 | –0.71 | 0.27 |
| diff. from species elevation midpoint | 0.05 | –0.29 | 0.39 | –0.03 | –0.67 | 0.65 |
| breeding status (nonbreeding) | **0.36** | **0.00** | **0.74** | **–0.83** | **–1.60** | **–0.11** |
| season (rainy) | –0.20 | –0.53 | 0.11 | –0.49 | –1.13 | 0.14 |
| clutch size | –0.09 | –0.34 | 0.17 | –0.07 | –0.51 | 0.38 |
| diet (carnivore) | **0.09** | **0.00** | **0.18** | –0.06 | –0.23 | 0.11 |
| diet (herbivore) | *–0.11* | *–0.23* | *0.00* | –0.01 | –0.22 | 0.19 |
| diet (nectarivore) | –0.03 | –0.15 | 0.10 | 0.17 | –0.04 | 0.39 |

**Table S8:** Model summary of the alternative main model testing the association between baseline CORT and breeding status, breeding latitude, sex, migration, clutch size and diet. Model was controlled for species specific body mass and the IBMD, time from sunrise and blood sampling latency. Values with CrI95 not containing zero are highlighted in **bold** and regarded as significant support for an effect. Weakly supported trends in models are highlighted in *italic*. For the selected main model see Table 2 in the main text.

| | Baseline corticosterone | | |
|---|---|---|---|
| **predictor** | **b** | **LCrI** | **UCrI** |
| *ln species specific body mass* | *–0.18* | *–0.42* | *0.06* |
| IBMD | –0.02 | –0.07 | 0.02 |
| **blood sampling latency (min)** | **0.72** | **0.55** | **0.90** |
| **time since sunrise (h)** | **–0.05** | **–0.09** | **–0.01** |
| breeding status (nonbreeding) | 0.25 | –0.10 | 0.61 |
| **breeding latitude (tropical)** | **–0.70** | **–1.22** | **–0.20** |
| sex (female) | 0.04 | –0.18 | 0.25 |
| migration distance (1000km) | –0.02 | –0.17 | 0.13 |
| clutch size | –0.05 | –0.19 | 0.10 |
| **diet (carnivore)** | **0.11** | **0.03** | **0.20** |
| diet (nectarivore) | 0.06 | –0.05 | 0.17 |

**Table S9:** Model summary testing the association between CORT, sex and migration limited to temperate species only (n = 130 individuals of 44 species). Model was controlled for species specific body mass and the IBMD, time from sunrise and blood sampling latency. Values with CrI95 not containing zero are highlighted in **bold** and regarded as significant support for an effect. Weakly supported trends in models are highlighted in *italic*.

| | Baseline corticosterone | | |
|---|---|---|---|
| **predictor** | **b** | **LCrI** | **UCrI** |
| ln species specific body mass | 0.08 | –0.27 | 0.44 |
| IBMD | 0.00 | –0.07 | 0.07 |
| time from sunrise (h) | –0.01 | –0.07 | 0.06 |
| **blood sampling latency (min)** | **0.59** | **0.31** | **0.86** |
| sex (female) | –0.21 | –0.57 | 0.14 |
| migration distance (1000km) | 0.10 | –0.06 | 0.26 |

**Table S10:** Model summaries of elevational and seasonal (breeding status and rainy/dry season) variation in baseline CORT and TEST for tropical species only (except for *Emberiza gosling*; n = 169 individuals of 55 species). Models were controlled for species specific body mass and the IBMD, time from sunrise and blood sampling latency. Values with CrI95 not containing zero are highlighted in **bold** and regarded as significant support for an effect. Weakly supported trends in models are highlighted in *italic*.

| | Baseline corticosterone | | | Testosterone | | |
|---|---|---|---|---|---|---|
| **predictor** | **b** | **LCrI** | **UCrI** | **b** | **LCrI** | **UCrI** |
| ln species specific body mass | –0.26 | –0.62 | 0.12 | **–1.15** | **–1.81** | **–0.51** |
| IBMD | –0.02 | –0.08 | 0.03 | –0.08 | –0.22 | 0.06 |
| time from sunrise (h) | **–0.06** | **–0.11** | **–0.01** | –0.05 | –0.15 | 0.05 |
| blood sampling latency (min) | **0.86** | **0.62** | **1.09** | 0.2 | –0.25 | 0.66 |
| sex (female) | **0.26** | **0.00** | **0.51** | **–2.02** | **–2.64** | **–1.45** |
| species elevation midpoint (km a.s.l.) | –0.12 | –0.42 | 0.19 | –0.06 | –0.60 | 0.49 |
| elevational difference (km) | 0.10 | –0.26 | 0.46 | –0.02 | –0.70 | 0.69 |
| season (rainy) | *–0.26* | *–0.58* | *0.07* | –0.37 | –1.03 | 0.28 |
| breeding status (nonbreeding) | **0.47** | **0.09** | **0.85** | *–0.76* | *–1.59* | *0.01* |

**Table S11:** Model summaries of the model testing the association between stress induced CORT (available for 75 individuals of 24 species) and the magnitude of the stress reaction and baseline CORT (magnitude model only), breeding latitude, sex, migration, clutch size and diet. Model was controlled for species specific body mass and the IBMD, time from sunrise and stressor duration (time between first and second bleeding). Values with CrI95 not containing zero are highlighted in **bold** and regarded as significant support for an effect. Weakly supported trends in models are highlighted in *italic*.

| | Stress-induced CORT | | |
|---|---|---|---|
| **predictor** | **b** | **LCrI** | **UCrI** |
| ln species specific body mass | –0.02 | –0.54 | 0.48 |
| IBMD | 0.02 | –0.04 | 0.08 |
| time from sunrise (h) | –0.01 | –0.09 | 0.06 |
| stressor duration (min) | 1.91 | –4.52 | 8.36 |
| breeding latitude (tropical) | 0.07 | –1.11 | 1.24 |
| clutch size | 0.18 | –0.35 | 0.70 |
| sex (female) | –0.24 | –0.65 | 0.17 |
| **migration distance (1000km)** | **–0.35** | **–0.66** | **–0.04** |
| **ln baseline corticosterone** | **0.25** | **0.09** | **0.41** |

**Table S12:** Model summary of model testing the association between baseline blood glucose and baseline CORT (n = 300 individuals of 100 species), breeding latitude and sex. Model was controlled for species specific body mass and the IBMD, time from sunrise and blood sampling latency. Values with CrI95 not containing zero are highlighted in **bold** and regarded as significant support for an effect. Weakly supported trends in models are highlighted in *italic*.

| | **Baseline level of blood glucose** | | |
|---|---|---|---|
| **predictor** | **b** | **LCrI** | **UCrI** |
| **ln species specific body mass** | **–0.65** | **–1.23** | **–0.06** |
| IBMD | –0.02 | –0.09 | 0.06 |
| time from sunrise (h) | –0.03 | –0.10 | 0.03 |
| blood sampling latency (min) | 0.13 | –0.22 | 0.47 |
| **breeding latitude (tropical)** | **–0.79** | **–1.43** | **–0.15** |
| **sex (female)** | **0.52** | **0.17** | **0.87** |
| ln baseline corticosterone | –0.05 | –0.25 | 0.14 |

**Table S13:** Model summary of the alternative main model testing the association between TEST and breeding status, breeding latitude, sex, migration distance and clutch size as well as interaction between breeding latitude and sex. Model was controlled for species specific body mass and the IBMD, time from sunrise and blood sampling latency. Values with CrI95 not containing zero are highlighted in **bold** and regarded as significant support for an effect. Weakly supported trends in models are highlighted in *italic*. For the selected main model see Table 3 in the main text.

| | **Testosterone** | | |
|---|---|---|---|
| **predictor** | **b** | **LCrI** | **UCrI** |
| **ln species specific body mass** | **–0.87** | **–1.40** | **–0.34** |
| IBMD | –0.04 | –0.14 | 0.05 |
| *blood sampling latency (min)* | *0.33* | *–0.03* | *0.70* |
| time since sunrise (h) | –0.01 | –0.09 | 0.07 |
| **breeding status (nonbreeding)** | **–0.81** | **–1.65** | **0.00** |
| **breeding latitude (tropical)** | **–2.32** | **–3.50** | **–1.15** |
| **sex (female)** | **–2.27** | **–2.96** | **–1.60** |
| **migration distance (1000km)** | **–0.45** | **–0.78** | **–0.13** |
| *clutch size* | *–0.22* | *–0.54* | *0.09* |
| breeding latitude (tropical) × sex (female) | 0.19 | –0.72 | 1.11 |

#### Supplementary references

**Apfelbeck, B., Mortega, K.G., Flinks, H., Illera, J.C. & Helm, B.** 2017. Testosterone, territorial response, and song in seasonally breeding tropical and temperate stonechats. *BMC Evol. Biol.* **17**: 101. https://doi.org/10.1186/s12862-017-0944-9

**Bílková, Z., Adámková, M., Albrecht, T. & Šimek, Z.** 2019. Determination of testosterone and corticosterone in feathers using liquid chromatography-mass spectrometry. *J. Chromatogr. A* **1590**: 96–103. https://doi.org/10.1016/j.chroma.2018.12.069

**Billerman, S. M., Keeney, B. K., Rodewald, P. G. & Schulenberg T. S.** (eds.) 2022. *Birds of the World* Cornell Laboratory of Ornithology, Ithaca, NY, USA. https://birdsoftheworld.org/

**Bürkner, P.-C.** 2017. **brms**: An *R* Package for Bayesian Multilevel Models Using *Stan*. *J. Stat. Softw.* **80**. https://doi.org/10.18637/jss.v080.i01

**De Rosa, A., Castro, I. & Marsland, S.** 2022. The acoustic playback technique in avian fieldwork contexts: a systematic review and recommendations for best practice. *Ibis* **164**: 371–387. https://doi.org/10.1111/ibi.13033

**Deviche, P., Beouche-Helias, B., Davies, S., Gao, S., Lane, S. & Valle, S.** 2014. Regulation of plasma testosterone, corticosterone, and metabolites in response to stress, reproductive stage, and social challenges in a desert male songbird. *Gen. Comp. Endocrinol.* **203**: 120–131. https://doi.org/10.1016/j.ygcen.2014.01.010

**Deviche, P., Small, T., Sharp, P. & Tsutsui, K.** 2006. Control of luteinizing hormone and testosterone secretion in a flexibly breeding male passerine, the Rufous-winged Sparrow, *Aimophila carpalis*. *Gen. Comp. Endocrinol.* **149**: 226–235. https://doi.org/10.1016/j.ygcen.2006.06.004

**Gonzalez-Voyer, A., Thomas, G.H., Liker, A., Krüger, O., Komdeur, J. & Székely, T.** 2022. Sex roles in birds: Phylogenetic analyses of the influence of climate, life histories and social environment. *Ecol. Lett.* **25**: 647–660. https://doi.org/10.1111/ele.13938

**Mackworth-Praed, C.W. & Grant, C.H.B.** 1970. *Birds of West Central and Western Africa, volume 1*. Longman, London.

**Mersman, O., Trautman, H., Steuer, D., & Bornkamp, B**. (2023). truncnorm: Truncated Normal Distribution. https://CRAN.R-project.org/package=truncnorm

**R Core Team**. 2023. R: A Language and Environment for Statistical Computing. https://www.r-project.org

**Romero, L.M. & Reed, J.M.** 2005. Collecting baseline corticosterone samples in the field: is under 3 min good enough? *Comp. Biochem. Physiol. A. Mol. Integr. Physiol.* **140**: 73–79. https://doi.org/10.1016/j.cbpb.2004.11.004

**Schwabl, P., Bonaccorso, E. & Goymann, W.** 2016. Diurnal variation in corticosterone release among wild tropical forest birds. *Front. Zool.* **13**: 19. https://doi.org/10.1186/s12983-016-0151-3

**Shoari, N. & Dubé, J.-S.** 2018. Toward improved analysis of concentration data: Embracing nondetects. *Environ. Toxicol. Chem.* **37**: 643–656. https://doi.org/10.1002/etc.4046

**Therneau, T.M.** 2025. A Package for Survival Analysis in R. https://cran.r-project.org/package=survival

**Thieurmel, B. & Elmarhraoui, A.** 2017. suncalc: Compute Sun Position, Sunlight Phases, Moon Position and Lunar Phase. https://doi.org/10.32614/CRAN.package.suncalc
